# Molecular characterization of RIGI, TLR7 and TLR3 as immune response gene of indigenous ducks in response to Avian influenza

**DOI:** 10.1101/2020.09.28.316687

**Authors:** Aruna Pal, Abantika Pal, Pradyumna Baviskar

## Abstract

Avian influenza is an alarming disease, which has every possibility to evolve as human to human pandemic situation due to frequent mutation and genetic reassortment or recombination of Avian influenza(AI) virus. The greatest concern is that till date no satisfactory medicine or vaccines are available, leading to massive culling of poultry birds causing huge economic loss, and ban on export of chicken products, which emphasise the need develop alternative strategy for control of AI. In the current study we attempt to explore the molecular mechanism of innate immune potential of ducks against common viral diseases including Avian influenza. In the present study, we have characterized immune response molecules as duck TLR3, TLR7, and RIGI and predicted to have potent antiviral activities against different identified strains of Avian influenza through *in silico* studies (molecular docking). Future exploitation involve immunomodulation with the recombinant protein, transgenic or gene-edited chicken resistant to bird flu.

## Introduction

Ducks are observed to be very resistant to common poultry diseases, including viral disease compared to chicken^**1**^ and are commonly asymptomatic to Avian Influenza virus infection. There is clear lack of further systematic characterization of the indigenous ducks at the molecular level. In an effort to understand, we have studied this as a first step. Hence there is an urgent need to explore the innate immune response genes, particularly against viral infection.

Avian influenza is caused by single stranded RNA virus, negatively stranded which belongs to Orthomyxoviridae family^**2**^. It is commonly known as Bird Flu, since birds are the main host. Based on the antigenic differences of two surface proteins- Haemagglutinin and neuraminidase of Avian influenza virus have been mostly subtyped and nomenclature provided accordingly. Till date 18 subtypes of HA (H1-H18) and 11NA (N1-N11) have been detected^**3,4**^. H5, H7 and H9 were observed to be the most pathogenic subtypes of bird. Most of the H5, H7 subtypes were regarded as Highly pathogenic avian influenza (HPAI) virus, owing to the higher incidence and mortality of birds. The greatest concern is the lack of definite treatment or vaccination due to frequent mutation and reassortment of viral strain, regarded as antigenic shift and antigenic drift^**5,6**^. Due to massive culling of birds in affected area and ban on export of poultry products, WHO has regarded Avian influenza as one of the most economically effected zoonotic disease^**7**^.

The basic mechanism of host immunity against viral infection is generally different from other infectious agents such as bacteria, protozoa etc. Viruses utilize the host immune mechanism for its infection and further survival, thus allowing it to act as hijackers. Accordingly viruses employ the host cellular machinery for living normal cells through the process of invasion, multiplication within the host, in turn kill, damage, or change the cells and make the individual sick^8^.

As the virus gets an entry in the body, an immune response is triggered followed by local inflammatory signalling. Innate immune reaction is initially activated by conserved pathogen-associated molecular pattern (PAMPs), pattern recognition receptors (PRRs), retinoic acid-inducible gene (RIG)-I like receptors, MDA5, LGP 2, and toll-like receptor (TLRs) as TLR3, TLR7^9^. Viral nucleic acid binds to these receptors expressed on macrophages, microglia, dendritic cells, astrocytes and releases type-I interferon (IFN-I) and production of interferon-stimulated genes (ISGs)^10^. Interferon –I upregulate antiviral proteins and accordingly peripheral immune cells are stimulated and alter endothelial tight junction^11^. It has been observed that the absence of IFN_I signaling leads to prevention of microglial differentiation and decrease of peripheral myeloid cell patrolling^11^.

TLR7 is a member of the Toll-like receptor family, which recognizes single-stranded RNA in endosomes, which is a common feature of viral genomes^12^. TLR7 can recognize GU-rich single-stranded RNA. TRL7 was reported to have influences on viral infection in poultry and has been regarded as a vital component of antiviral immunity, particularly in ducks.^12^ RIG-I (retinoic acid-inducible gene I) or RIG-I like receptor dsRNA helicase enzyme is part of the RIG-I like receptor family, which also includes MDA5 and LGP2. These have been reported to function as a pattern recognition receptor that is a sensor for viruses such as influenza A and others as Sendai virus, and flavivirus^13^. RIG-I typically recognizes short 5′ triphosphate uncapped double-stranded or single-stranded RNA^14^. RIG-I and MDA5 are the viral receptors, acting through a common adapter MAVS and triggers an antiviral response through type-I interferon response. RIG1 is an important gene conferring antiviral immunity for ducks, particularly avian influenza^13^.

TLR3 is another member of the toll-like receptor (TLR) family. Infectious agents express PAMP (Pathogen associated molecular patterns), which is readily recognized by TLR3, which in turn secretes cytokines responsible for effective immunity. It recognizes dsRNA associated with a viral infection, and induces the activation of IRF3, unlike all other toll-like receptors which activate NF-κB^15^. IRF3 ultimately induces the production of type I interferons, which is ultimately responsible for host defense against viruses^16^. In our lab, earlier we had studied immunogenetics against bacterial disease with identified immune response molecule as CD14 gene in goat ^**17, 18**^, cattle ^**19**^, buffalo ^**20, 21**^. We reported for the first time the role mitochondrial cytochrome B gene for immunity in sheep^**23**^ and immune-response genes in Haringhata Black Chicken ^**24**^.

Indigenous duck population in the Indian subcontinent was observed to have better immunity against viral infections and so far, no systematic studies were undertaken. Thus, the present study was conducted with the aim of molecular characterization of immune response genes (TLR3, TLR7, RIGI) of duck, provide initial Proteomics study and prediction of binding site with multiple strains of Avian influenza virus through *in silico* studies (molecular docking) and establishment of disease-resistant genes of ducks through Quantitative PCR.

## Materials and methods

### Animals, Sample Collection, RNA Isolation

#### Birds

Duck samples were collected from the different agro-climatic region of West Bengal, India from farmer’s herd. The chicken breeds as Haringhata Black, Aseel were maintained in the university farm (West Bengal University of Animal and Fishery Sciences). Samples from other poultry species as Guineafowl, goose were also collected from the university farm. Samples from Turkey and quail were collected from State Poultry farm, Animal Resource Development Dept, Tollygunge, Govt. of West Bengal, India. The birds were vaccinated against routine diseases as Ranikhet disease, fowl pox. Six male birds (aged 4-5 months) were considered under each group for this study and are maintained under uniform managemental conditions.

All experiments were conducted in accordance with relevant guidelines and regulations of Institutional Animal Ethics committee and all experimental protocols were approved by the Institutional Biosafety Committee, West Bengal University of Animal and Fishery Sciences, Kolkata.

The total RNA was isolated from the ileocaecal junction of Duck, Haringhata Black chicken, Aseel and other poultry species as Guineafowl, goose, using Ribopure kit (Invitrogen), following manufacturer’s instructions and was further used for cDNA synthesis^17, 23^.

### Materials

Taq DNA polymerase, 10X buffer, dNTP were purchased from Invitrogen, SYBR Green qPCR Master Mix (2X) was obtained from Thermo Fisher Scientific Inc. (PA, USA). L-Glutamine (Glutamax 100x) was purchased from Invitrogen corp., (Carlsbad, CA, USA). Penicillin-G and streptomycin were obtained from Amresco (Solon, OH, USA). Filters (Millex GV. 0.22 μm) were purchased from Millipore Pvt. Ltd., (Billerica, MA, USA). All other reagents were of analytical grade.

### Synthesis, Confirmation of cDNA and PCR Amplification of TLR3, RIGI and TLR7 gene

The 20□μL reaction mixture contained 5□μg of total RNA, 0.5□μg of oligo dT primer (16–18□mer), 40□U of Ribonuclease inhibitor, 10□M of dNTP mix, 10□mM of DTT, and 5□U of MuMLV reverse transcriptase in reverse transcriptase buffer. The reaction mixture was gently mixed and incubated at 37°C for 1 hour. The reaction was stopped by heating the mixture at 70°C for 10 minutes and chilled on ice. The integrity of the cDNA was checked by PCR. To amplify the full-length open reading frame (ORF) of gene sequence, a specific primers pair was designed based on the mRNA sequences of Gallus gallus by DNASTAR software. The primers have been listed in Table 1. 25□μL reaction mixture contained 80–100□ng cDNA, 3.0□μL 10X PCR assay buffer, 0.5□μL of 10□mM dNTP, 1□U Taq DNA polymerase, 60□ng of each primer, and 2□mM MgCl2. PCR-reactions were carried out in a thermocycler (PTC-200, MJ Research, USA) with cycling conditions as, initial denaturation at 94°C for 3□min, denaturation at 94°C for 30□sec, varying annealing temperature (as mentioned in Table 1) for 35□sec, and extension at 72°C for 3□min was carried out for 35 cycles followed by final extension at 72°C for 10□min.

**Table 1:**
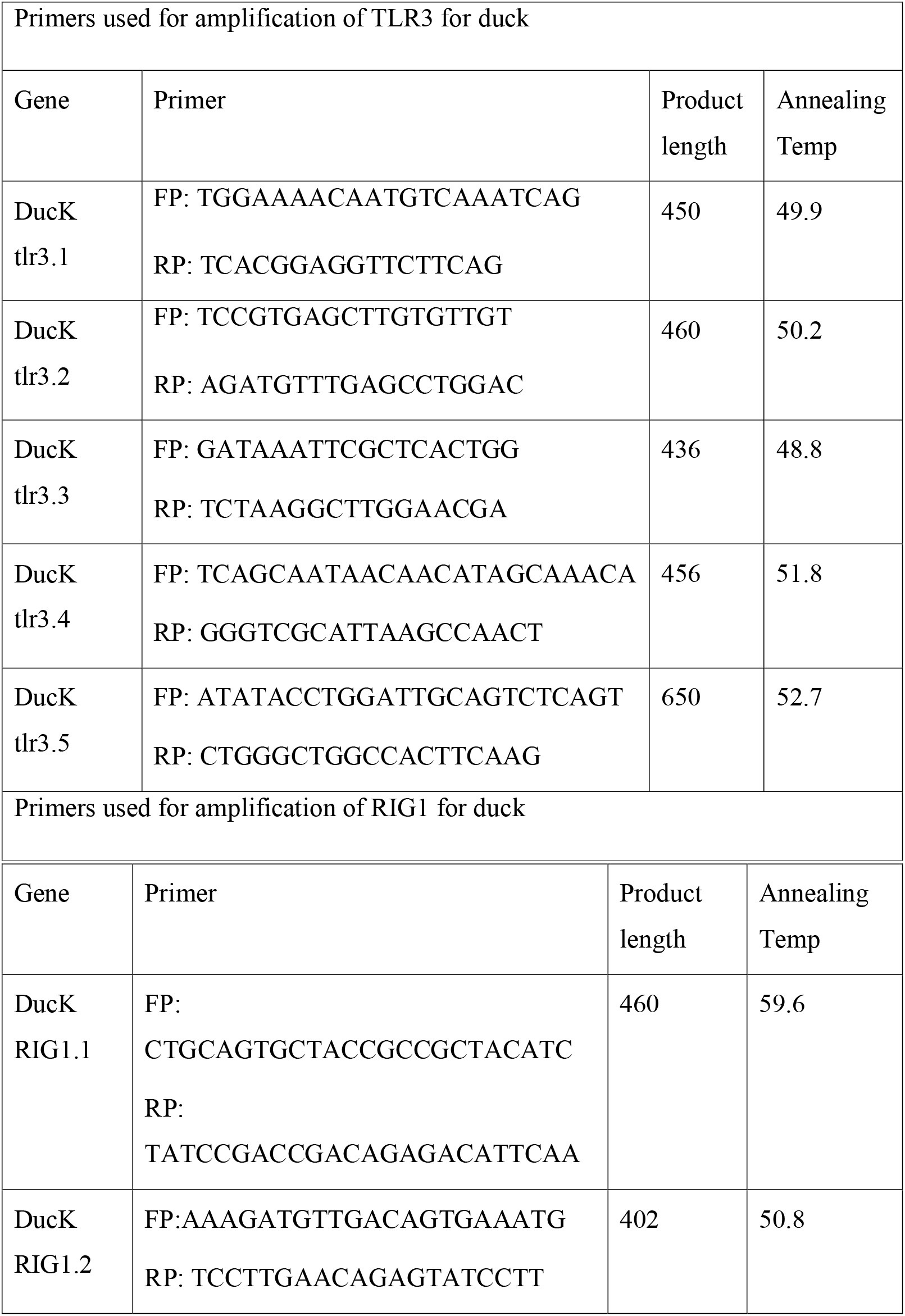
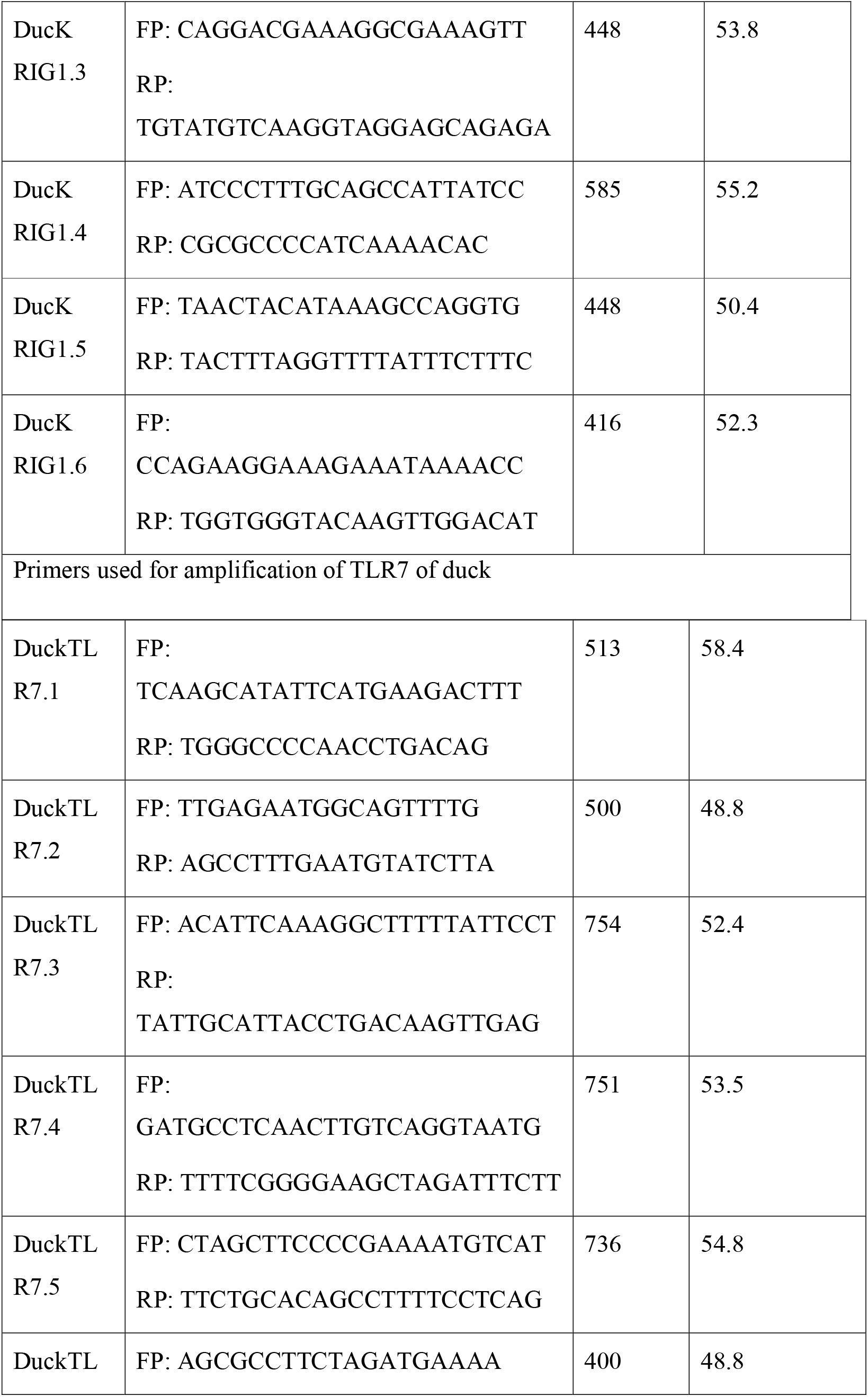
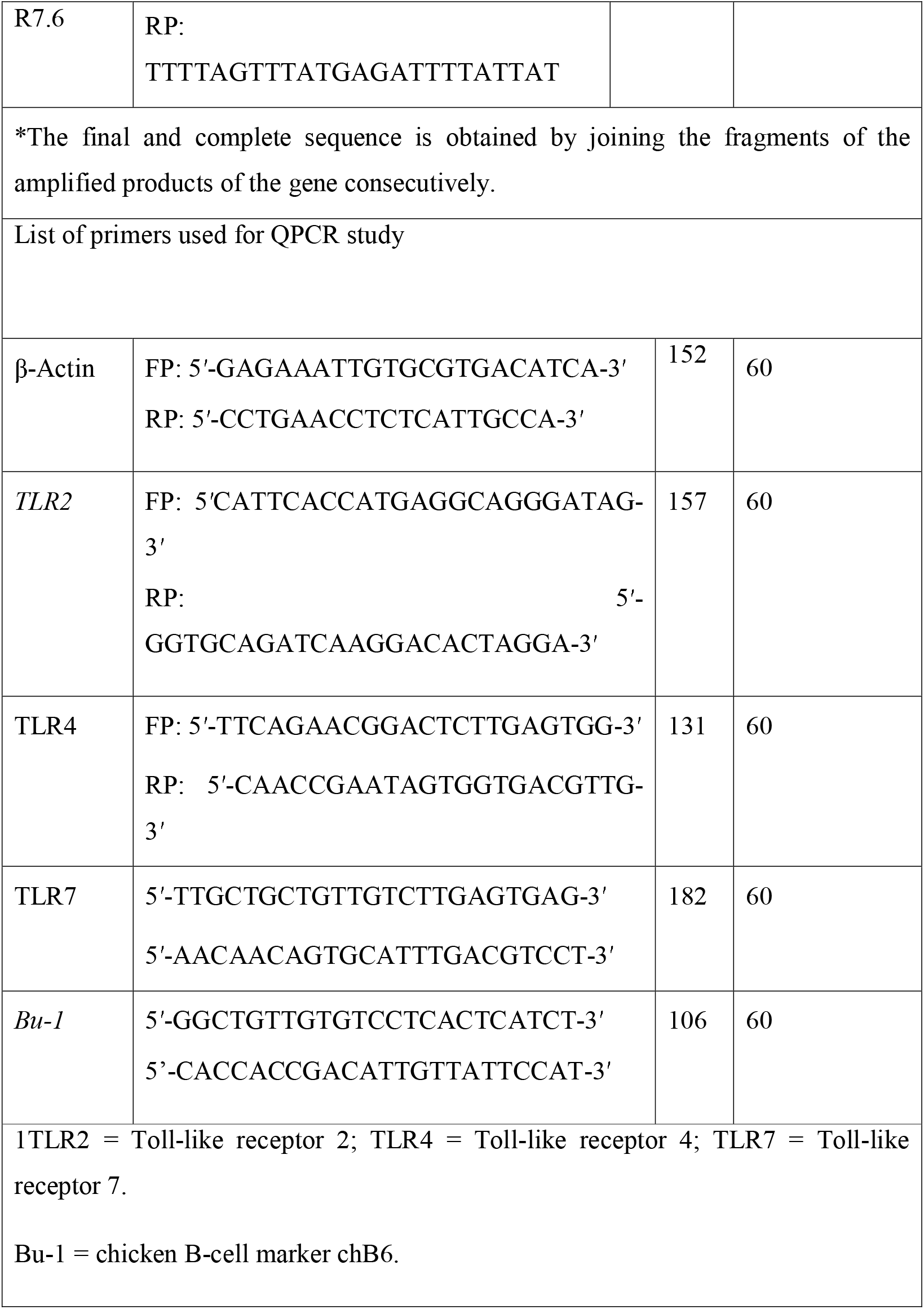
List of primers used for amplification of TLR3, RIG1 and TLR7 genes in indigenous duck.

### cDNA Cloning and Sequencing

PCR amplicons verified by 1% agarose gel electrophoresis were purified from gel using Gel extraction kit (Qiagen GmbH, Hilden, Germany) and ligated into a pGEM-T easy cloning vector (Promega, Madison, WI, USA) following manufacturers’ instructions. The 10□μL of the ligated product was directly added to 200□μL competent cells, and heat shock was given at 42°C for 45□sec. in a water bath, and cells were then immediately transferred on chilled ice for 5□min., and SOC was added. The bacterial culture was pelleted and plated on LB agar plate containing Ampicillin (100□mg/mL) added to agar plate @ 1□:□1000, IPTG (200□mg/mL) and X-Gal (20□mg/mL) for blue-white screening. Plasmid isolation from overnight-grown culture was done by small-scale alkaline lysis method. Recombinant plasmids were characterized by PCR using gene-specific primers and restriction enzyme digestion based on reported nucleotide sequence for cattle. The enzyme EcoR I (MBI Fermentas, USA) is used for fragment release. Gene fragment insert in the recombinant plasmid was sequenced by an automated sequencer (ABI prism) using the dideoxy chain termination method with T7 and SP6 primers (Chromous Biotech, Bangalore).

### Sequence Analysis

The nucleotide sequence so obtained was analyzed for protein translation, sequence alignments, and contigs comparisons by DNASTAR Version 4.0, Inc., USA. The novel sequence was submitted to the NCBI Genbank and accession number was obtained which is available in public domain now.

### Study of Predicted TLR3, TLR7 and RIG1 peptide Using Bioinformatics Tools

The predicted peptide sequence of TLR3, TLR7, and RIG1 of indigenous duck was derived by Edit sequence (Lasergene Software, DNASTAR) and then aligned with the peptide of other chicken breed and avian species using Megalign sequence Programme of Lasergene Software (DNASTAR). Prediction of the signal peptide of the CD14 gene was conducted using the software (Signal P 3.0 Sewer-prediction results, Technical University of Denmark). Estimation of Leucine percentage was conducted through manually from the predicted peptide sequence. Di-sulfide bonds were predicted using suitable software (http://bioinformatics.bc.edu/clotelab/DiANNA/) and by homology search with other species.

Protein sequence-level analysis study was carried out with specific software (http://www.expasy.org./tools/blast/) for determination of leucine-rich repeats (LRR), leucine zipper, N-linked glycosylation sites, detection of Leucine-rich nuclear export signals (NES), and detection of the position of GPI anchor. Detection of Leucine-rich nuclear export signals (NES) was carried out with NetNES 1.1 Server, Technical University of Denmark. Analysis of O-linked glycosylation sites was carried out using NetOGlyc 3.1 server (http://www.expassy.org/), whereas the N-linked glycosylation site was detected by NetNGlyc 1.0 software (http://www.expassy.org/). Detection of Leucine-zipper was conducted through Expassy software, Technical University of Denmark ^24^. Regions for alpha-helix and beta-sheet were predicted using NetSurfP-Protein Surface Accessibility and Secondary Structure Predictions, Technical University of Denmark^25^. Domain linker prediction was done according to the software developed^26^. LPS-binding site^27^, as well as LPS-signaling sites^28^, were predicted based on homology studies with other species polypeptide.

### Three-dimensional structure prediction and Model quality assessment

The templates which possessed the highest sequence identity with our target template were identified by using PSI-BLAST (http://blast.ncbi.nlm.nih.gov/Blast). The homology modeling was used to build a 3D structure based on homologous template structures using PHYRE2 server^29^. The 3D structures were visualized by PyMOL (http://www.pymol.org/) which is an open-source molecular visualization tool. Subsequently, the mutant model was generated using PyMoL tool. The Swiss PDB Viewer was employed for controlling energy minimization. The structural evaluation along with a stereochemical quality assessment of predicted model was carried out by using the SAVES (Structural Analysis and Verification Server), which is an integrated server (http://nihserver.mbi.ucla.edu/SAVES/). The ProSA (Protein Structure Analysis) webserver (https://prosa.services.came.sbg.ac.at/prosa) was used for refinement and validation of protein structure^30^. The ProSA was used for checking model structural quality with potential errors and the program shows a plot of its residue energies and Z-scores which determine the overall quality of the model. The solvent accessibility surface area of the IR genes was generated by using NetSurfP server (http://www.cbs.dtu.dk/services/NetSurfP/)^31^. It calculates relative surface accessibility, Z-fit score, the probability for Alpha-Helix, probability for beta-strand and coil score, etc. TM align software was used for the alignment of 3 D structure of IR protein for different species and RMSD estimation to assess the structural differentiation ^32^. The I-mutant analysis was conducted for mutations detected to assess the thermodynamic stability^33^. Provean analysis was conducted to assess the deleterious nature of the mutant amino acid^34^.

### Molecular Docking

Molecular docking is a bioinformatics tool used for *in silico* analysis for the prediction of the binding mode of a ligand with a protein 3D structure. Patch dock is an algorithm for molecular docking based on shape complementarity principle ^35^. Patch Dock algorithm was used to predict ligand-protein docking for surface antigen for Avian influenza (H-antigen and NA antigen) with the molecules for innate immunity against viral infection as TLR3, TLR7, and RIG1. Firedock were employed for further confirmation^**36**^. The amino acid sequence for the surface antigen (Haemagglutinin and neuraminidase) from different strains of Avian influenza were retrieved from gene bank. Haemagglutinin segment 4 sequence were collected from Indian sub-continent as H5N1(Acc no. KR021385, Protein id. AKD00332), H4N6 (Acc no.JX310059, Protein id. AF082958), H6N2 (Acc no. KU598235, Protein id. AMH93683), H9N2 (Acc no. 218091, Protein id. AAG53040). Neuraminidase segment 6 were collected from Indian sub continent as H5N1(Acc no. KT867346, Protein id. ALK80150), H4N6 (Acc no. JX310060, Protein id. AF082959), H6N2 (Acc no. KU598237, Protein id. AMH93685) to be employed as Ligand. The receptor molecule employed were TLR3 (Gene bank accession number KX865107, NCBI) and derived protein as ASW23003), RIGI (Gene bank accession number KX865107, protein ASW23002 from NCBI) and TLR7 (Gene bank Accession no.MK986726, NCBI) for duck sequenced and characterized in our lab.

### Assessment of antigenic variability among different strains of Avian influenza

MAFFT software^**37**^ was employed for detection of amino acid variability and construction of phylogenetic tree for different strains of Avian influenza detected in duck in Indian sub continent.

The amino acid sequence for the surface antigen (Haemagglutinin and neuraminidase) from different strains of Avian influenza were retrieved from gene bank. Haemagglutinin segment 4 sequence were collected from Indian sub-continent as H5N1(Acc no. KR021385, Protein id. AKD00332), H4N6 (Acc no.JX310059, Protein id. AF082958), H6N2 (Acc no. KU598235, Protein id. AMH93683), H9N2 (Acc no. 218091, Protein id. AAG53040).

Neuraminidase segment 6 were collected from Indian sub continent as H5N1(Acc no. KT867346, Protein id. ALK80150), H4N6 (Acc no. JX310060, Protein id. AF082959), H6N2 (Acc no. KU598237, Protein id. AMH93685.1)

### Protein-protein interaction network depiction

In order to understand the network of TLR3, TLR7 and RIG1 peptide, we performed analysis with submitting FASTA sequences to STRING 9.1^38^. Confidence scoring was used for functional analysis. Interactions with score < 0.3 are considered as low confidence, scores ranging from 0.3 to 0.7 are classified as medium confidence and scores > 0.7 yield high confidence. The functional partners were depicted. KEGG analysis also depicts the functional association of TLR3, TLR7 and RIG1 peptide with other related proteins (KEGG: Kyoto Encyclopedia of Genes and Genomes – GenomeNet, https://www.genome.jp/kegg/)

### Real-Time PCR (qRT-PCR)

An equal amount of RNA (quantified by Qubit fluorometer, Invitrogen), wherever applicable, were used for cDNA preparation (Superscript III cDNA synthesis kit; Invitrogen). All qRT-PCR reactions were conducted on the ABI 7500 fast system. Each reaction consisted of 2 μl cDNA template, 5 μl of 2X SYBR Green PCR Master Mix, 0.25 μl each of forward and reverse primers (10 pmol/μl) and nuclease-free water for a final volume of 10 μl. Each sample was run in duplicate. Analysis of real-time PCR (qRT-PCR) was performed by delta-delta-Ct (^ΔΔ^Ct) method. The primers used for QPCR analysis have been listed as per Table 1.

### Comparison of TLR3, TLR7 and RIG1 structure of indigenous ducks with respect to chicken

Nucleotide variation for the proteins was detected from their nucleotide sequencing and amino acid variations were estimated (DNASTAR). 3D structure of the derived protein was estimated for both indigenous ducks and chicken by Pymol software. PDB structure of the respective proteins was derived from *PHYRE* software^39^. We also employed *Modeller software* for protein structural modelling^**40**^ for better confirmation. Alignment of the structure of TLR3, TLR7 and RIGI duck with Chicken was conducted by TM Align software^**41**^.

## Results

### Molecular characterization of TLR3 gene

A toll-like receptor are a group of pattern recognition receptor effective against a wide range of pathogens. TLR3 gene of indigenous ducks have been characterized with 2688 bp Nucleotide (Gene bank accession number KX865107, NCBI) and derived protein as ASW23003.1. The 3D protein structure (Fig 1a) with surface view (Fig 1b) has been depicted, Helix Light blue, sheet red, loop pink, Blue spheres as Disulphide bonds.

**Fig1 a:**
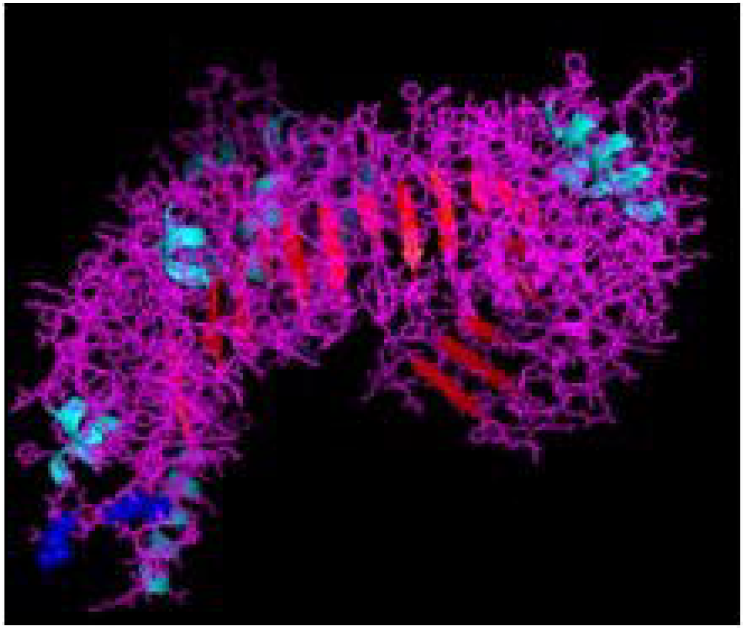
TLR3 molecule of duck (Secondary structure with disulphide bond) as blue sphere

**Fig1b:**
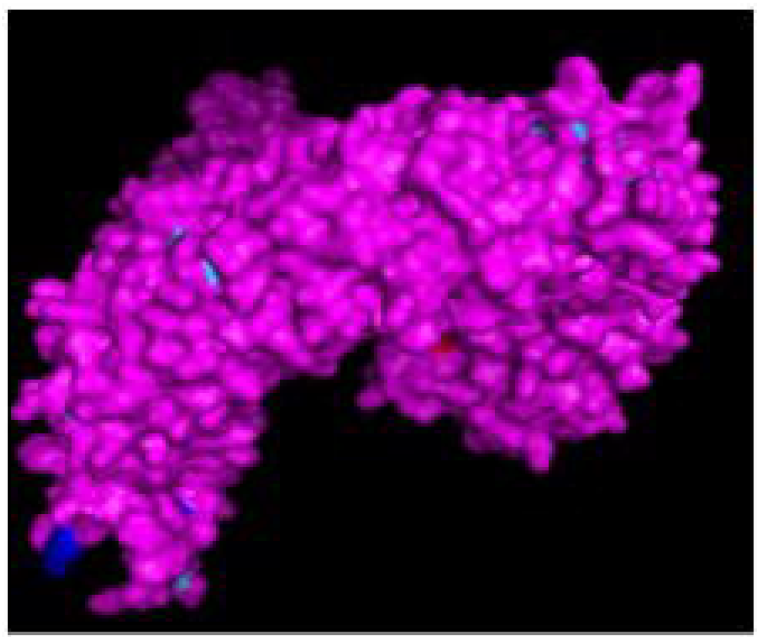
TLR3 molecule of duck (Secondary structure with disulphide bond) surface view

Post-translational modification sites for TLR3 of duck have been depicted in Fig 1c to 1e. Fig 1c reveals 3D structure of TLR3 of duck with the sites for leucine zipper (151-172 amino acid position, yellow surface), GPI anchor (aa position 879, red sphere), leucine-rich nuclear export signal (aa position 75-83, blue sphere), LRRNT (aa position 37-51, magenta sphere), LRRCT (aa position 664-687, orange sphere). Fig 1d depicts the 3D structure of TLR3 of duck with the site for TIR (amino acid position 748-890, blue sphere) and the sites for leucine-rich receptor-like protein kinase (amino acid position 314-637, orange mesh). Fig 1e represents the sites for leucine-rich repeats as spheres at aa sites 53-74 (blue), 77-98 (red), 101-122 (yellow-orange), 125-145 (hot pink), 148-168 (cyan), 172-195 (orange), 198-219 (grey), 275-296 (raspberry), 299-319 (split pea), 346-367 (purple-blue), 370-393 (sand), 422-444 (violet), 447-468 (deep teal), 497-518 (olive), 521-542 (green), 553-574 (grey), 577-598 (salmon), 601-622 (density). The other sites for post-translational modification as observed were 16 sites for N-linked glycosylation, 8 sites for casein kinase 2 phosphorylation, 8 sites for myristylation, 9 sites for phosphokinase phosphorylation. In comparison for TLR3 among the avian species, 51 amino acid variations were observed, which contribute to various important domains of TLR3. These domains are LRR, LRRCT, TIR (Table 2).

**Fig 1c:**
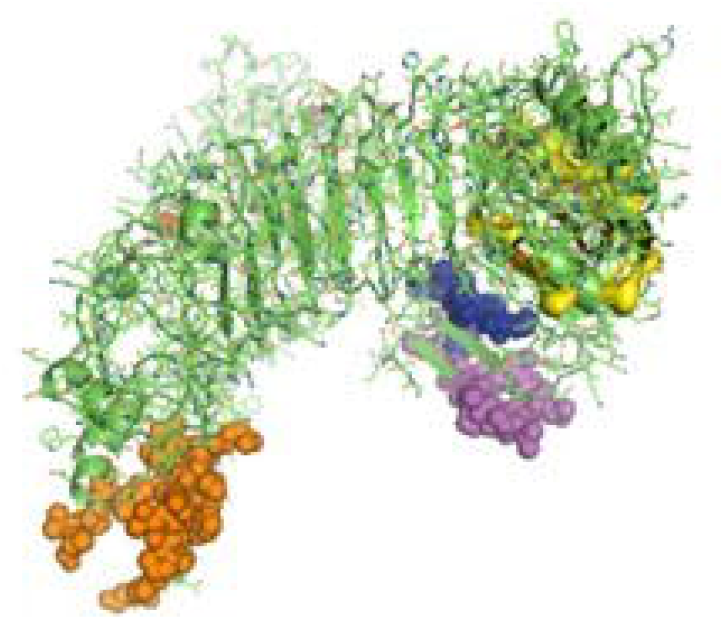
3D structure of TLR3 of duck: Yellow surface: Leucine zipper (151-172), Red sphere: GPI anchor (879), Blue sphere: Leucine rich nuclear export signal, Magenta sphere: LRRNT(37-51), Orange sphere: LRRCT (664-687)

**Fig 1d:**
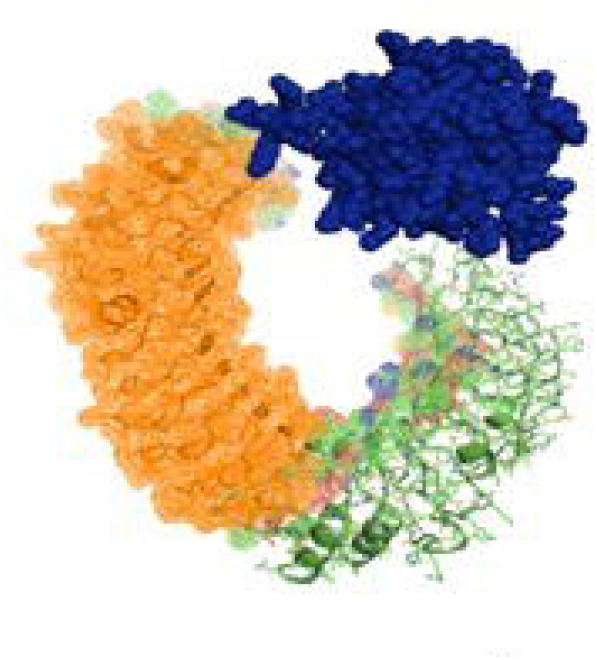
3D structure of TLR3 of indigenous duck Blue sphere:TIR(748-890), Orange mesh: Leucine rich receptor like proteinkinase

**Fig 1e:**
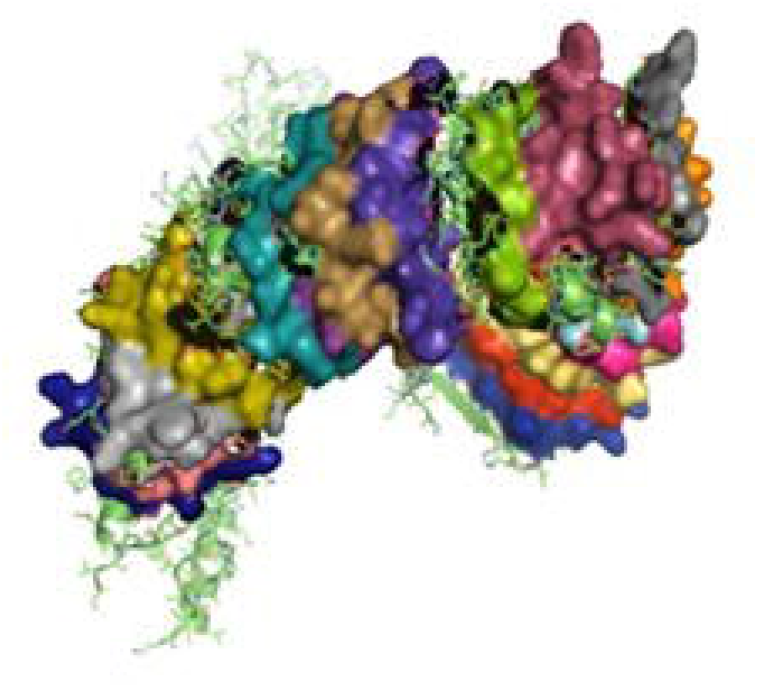
3D structure of TLR3 of duck with Leucine rich repeat

**Table 2:** Amino acid variations for TLR3 gene in Duck with other poultry species

| Sl no. | Position | Duck | Chicken | Turkey | Goose | Domain |
| --- | --- | --- | --- | --- | --- | --- |
| 1 | 42 | K | E | K | K | LRRNT |
| 2 | 61 | H | L | H | H | LRR1 |
| 3 | 68 | C | V | V | C | LRR1 |
| 4 | 69 | H | P | P | P | LRR1 |
| 5 | 70 | A | E | E | A | LRR1 |
| 6 | 74 | R | Q | E | K | LRR1 |
| 7 | 77 | K | N | N | K | LRR2 |
| 8 | 92 | Q | K | Q | Q | LRR2 |
| 9 | 94 | E | O | E | E | LRR2 |
| 10 | 106 | V | K | K | V | LRR3 |
| 11 | 119 | A | V | V | T | LRR3 |
| 12 | 137 | D | E | E | D | LRR4 |
| 13 | 166 | L | L | L | W | LRR5 |
| 14 | 179 | C | Y | Y | C | LLR6 |
| 15 | 187 | K | N | K | K | LLR6 |
| 16 | 192 | S | K | K | S | LLR6 |
| 17 | 200 | N | N | N | K | LLR7 |
| 18 | 212 | F | V | F | F | LRR7 |
| 19 | 213 | H | Q | H | H | LRR7 |
| 20 | 285 | Y | S | S | S | LRR8 |
| 21 | 299 | N | K | N | N | LRR9 |
| 22 | 306 | K | E | E | K | LRR9 |
| 24 | 310 | S | I | I | I | LRR9 |
| 28 | 317 | S | L | L | S | LRR9 |
| 29 | 319 | Y | Y | Y | H | LRR9 |
| 30 | 346 | Y | H | Y | Y | LRR10 |
| 31 | 355 | N | N | H | N | LRR10 |
| 32 | 360 | R | R | Q | R | LRR10 |
| 33 | 370 | N | K | K | N | LRR11 |
| 34 | 378 | S | Y | Y | S | LRR11 |
| 35 | 382 | I | T | I | I | LRR11 |
| 36 | 390 | T | K | K | T | LRR11 |
| 37 | 423 | H | Q | Q | H | LRR12 |
| 38 | 435 | S | N | N | S | LRR12 |
| 39 | 444 | K | E | E | K | LRR12 |
| 40 | 468 | S | I | I | S | LRR13 |
| 41 | 497 | Q | R | R | Q | LRR14 |
| 42 | 521 | H | H | Y | H | LRR15 |
| 43 | 522 | K | E | K | K | LRR15 |
| 44 | 539 | H | C | Q | H | LRR15 |
| 45 | 571 | Q | H | H | Q | LRR |
| 46 | 577 | F | H | Q | F | LRR |
| 47 | 581 | Y | D | N | Y | LRR |
| 48 | 601 | T | T | N | T | LRR |
| 49 | 619 | E | N | D | V | LRR |
| 50 | 686 | V | A | A | A | LRRCT |
| 51 | 766 | I | T | T | I | TIR1 |

### Molecular characterization of RIGI of duck

RIGI of duck have been characterized (Gene bank accession number KX865107, protein ASW23002 from NCBI). 3 D structure of RIGI of duck is depicted in Fig 2a and Fig 2b (surface view). RIGI is an important gene conferring antiviral immunity. A series of post-translational modification and various domains for its important function have been represented.

**Fig 2a:**
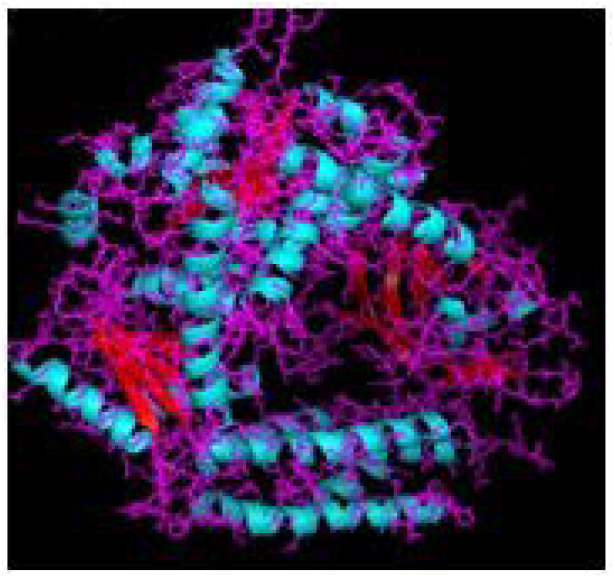
3D structure of RIG1 of duck

**Fig 2b:**
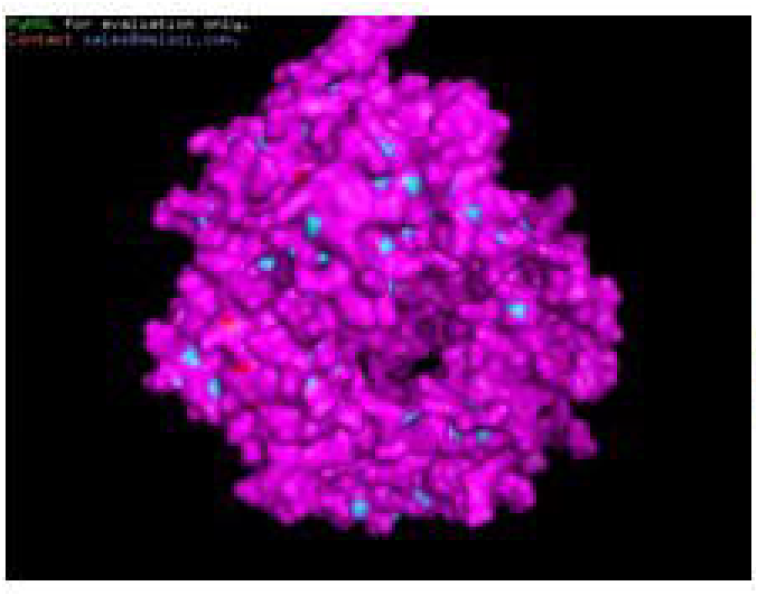
3D structure of RIG1 of duck, surface view

**Fig 2c:**
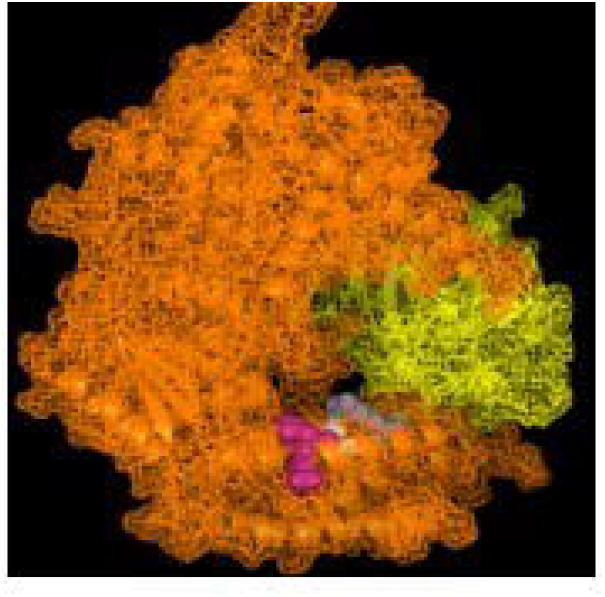
3D structure of RIG1 of duck with sites for helicase insert domain, helicase domain interface (polypeptide binding)

CARD_RIG1 (Caspase activation and recruitment domain found in RIG1) have been depicted in amino acid position 2-91, 99-188. CARD2 interaction site (17-20, 23-24, 49-50, 79-84), CARD1 interface (100, 103, 130-135, 155, 159,161-162), helical insert domain interface (101, 104-105, 107-108, 110-112, 114-115, 139, 143-145, 147-148, 151, 180, 183-184, 186 aa) have been depicted at RIG1 of duck. Fig 3c depicts helicase insert domain (242-800 aa) as orange mesh, helicase domain interface (polypeptide binding) as (511-512aa warm pink, 515aa white, 519aa grey). Fig 2d depicts double-stranded RNA binding site (nucleotide-binding) at amino acid positions 832 (red), 855 (green), 876-877 (blue), 889-891 (magenta), 911 (white). The sites for RD interface (polypeptide binding) and RIG-I-C (C terminal domain of retinoic acid-inducible gene, RIG-I protein, a cytoplasmic viral RNA receptor) have been depicted in Fig 2e. The site for RIG-I-C as amino acid position 807-921 represented by a mesh of pale green tints. The sites for RD interface have been depicted as amino acid positions 519 (red sphere), 522-523 (magenta), 536-537 (orange), 540 (grey).

**Fig 2d:**
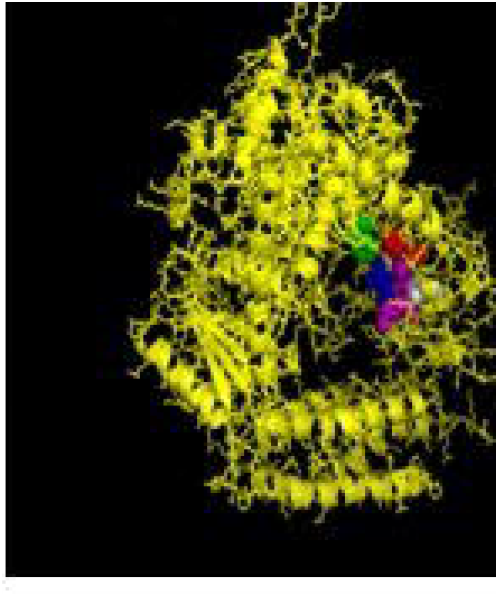
3D structure of RIG1 of duck depicting dsRNA binding site

**Fig 2e:**
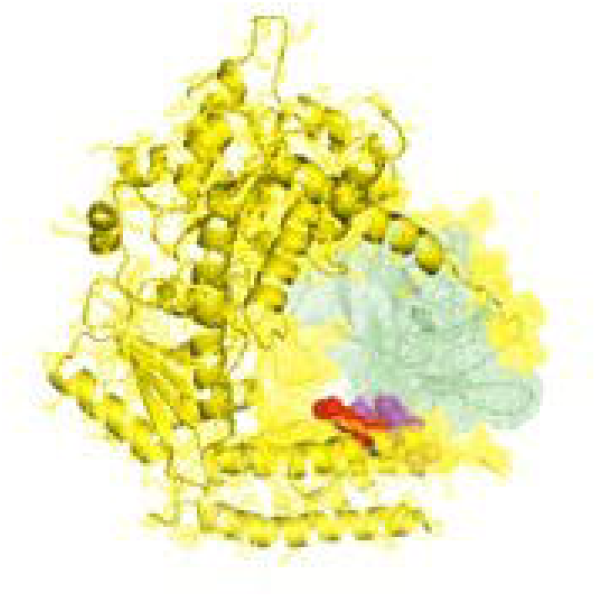
3D structure of RIGI of duck with RD interface and RIG IC

Fig 2f depicts the sites for the zinc-binding domain of RIG1 of duck as amino acid positions 812 (firebrick), 815 (marine blue), 866 (green), 871 (hot pink). The sites for RNA binding have been depicted as 511-512 (hot pink), 515 (cyan-deep teal), 519 (grey) in Fig 2g.

**Fig 2f:**
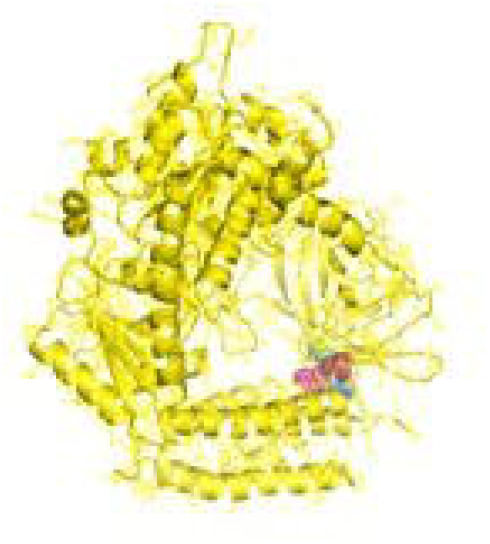
3D structure of RIG1 of duck for Zinc binding site

**Fig 2g:**
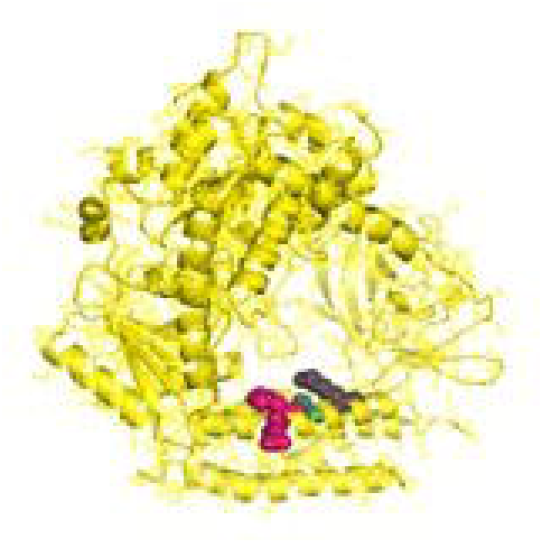
3D structure of RIG1 of duck of RNA binding site

### Molecular characterization of TLR7 of duck

TLR7 gene has been characterized in duck (Gene bank Accession no.MK986726, NCBI). The 3D structure of TLR7 is depicted in Fig 3a and Fig 3b(surface view). TLR7 is rich in leucine-rich repeat (LRR) as depicted in Fig 3c. The LRR sites are 104-125 (red sphere), 166-187(green), 188-210 (blue), 243-264(yellow), 265-285(magenta), 288-309(cyan), 328-400(orange), 435-455, 459-480(gray), 534-555(warm pink), 558-628(split pea), 691-712(purple-blue), 715-762(sand), 764-824(deep teal).

**Fig 3a:**
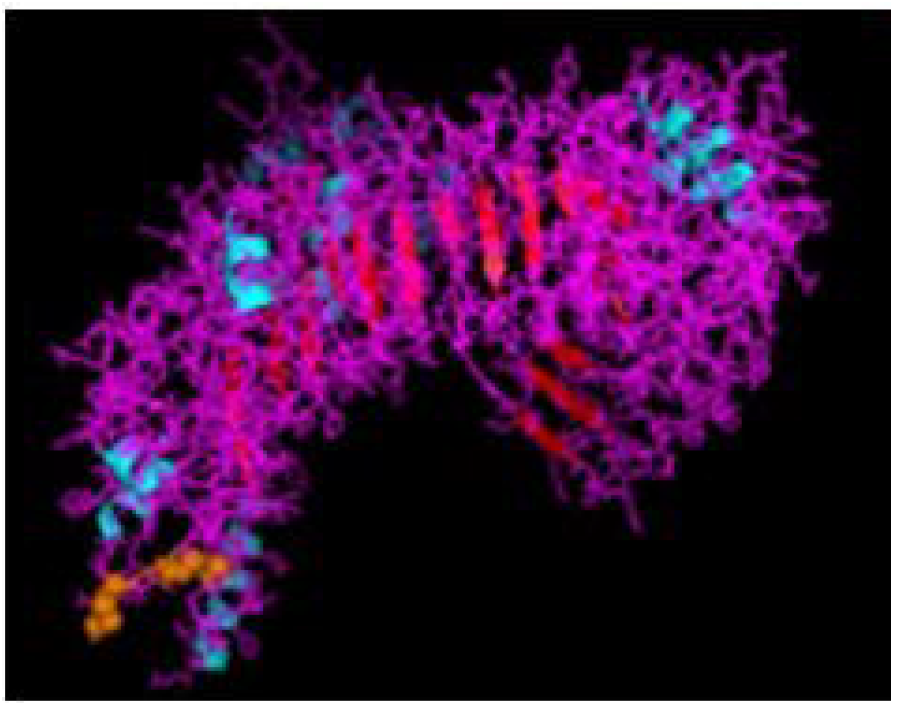
3D structure of TLR7 of indigenous Duck

**Fig 3b:**
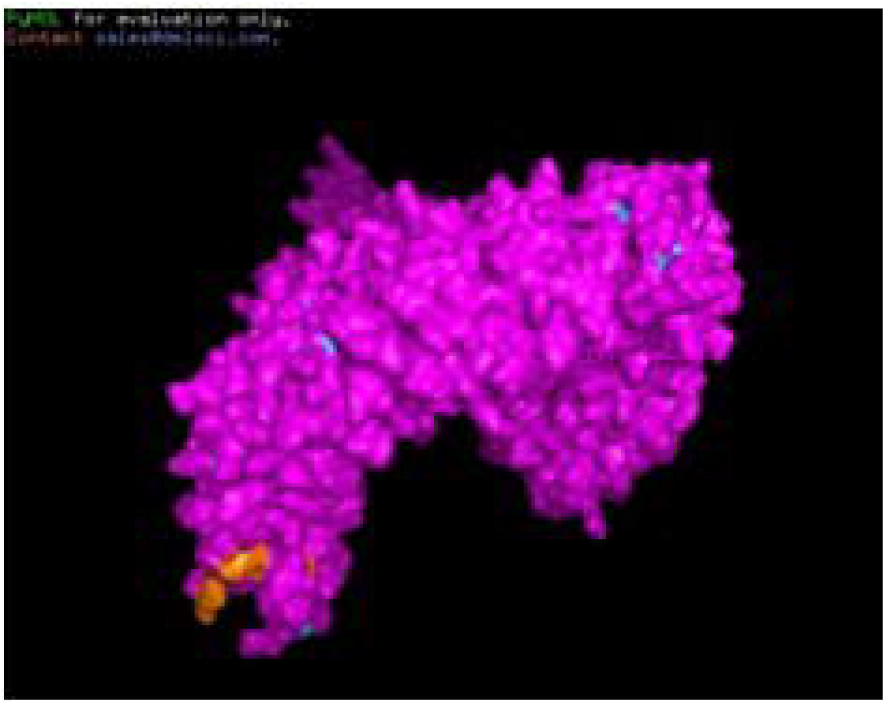
3D structure of TLR7 of Duck (surface view)

**Fig 3c:**
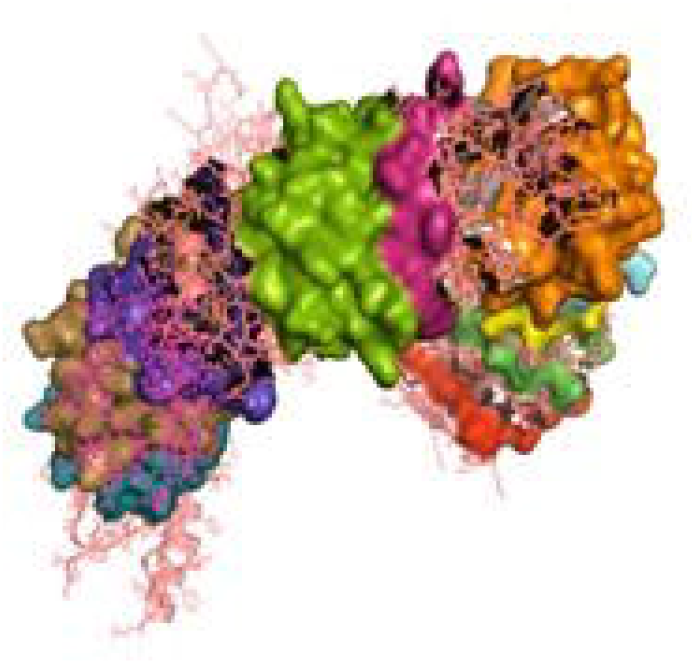
3D structure of TLR7 of duck with LRR region

The other domains are GPI anchor at 1072 amino acid position (red sphere), domain linker sites as 294-317(green sphere), 467-493 (split pea sphere) (Fig 3d). TIR site had been identified as 929-1076 amino acid position (blue sphere) and cysteine-rich flanking region, C-terminal as 823-874 amino acid position (hot pink) as depicted. The site for LRRNT (Leucine-rich repeat N-terminal domain) of TLR7 had been identified at amino acid position 75-107(red surface), TPKR-C2 (Tyrosine-protein kinase receptor C2 Ig like domain) at amino acid position 823-869 (blue surface) and GPI anchor as a green sphere (Fig 3e). Fig 3f represents the transmembrane site for TLR7 of duck.

**Fig 3d:**
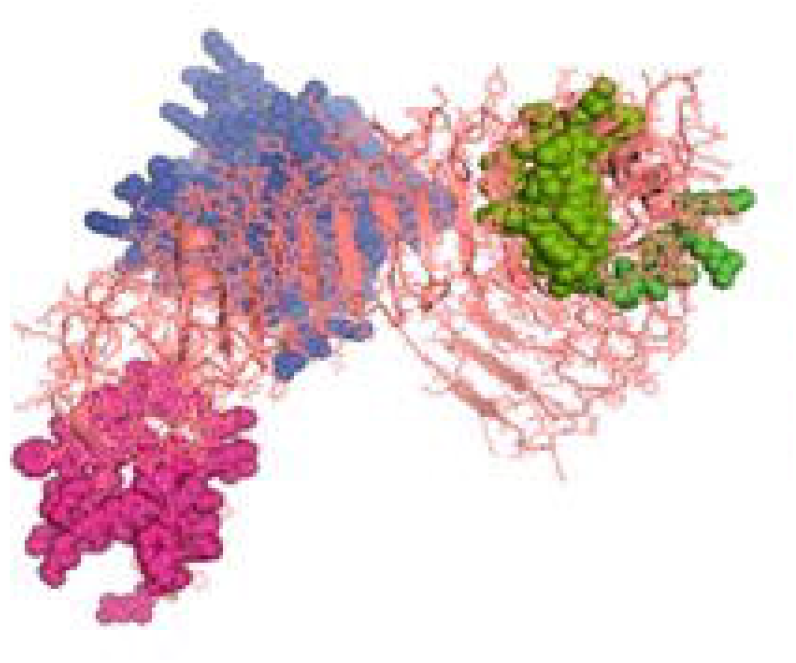
3D structure for TLR7 of duck-Domains

**Fig 3e:**
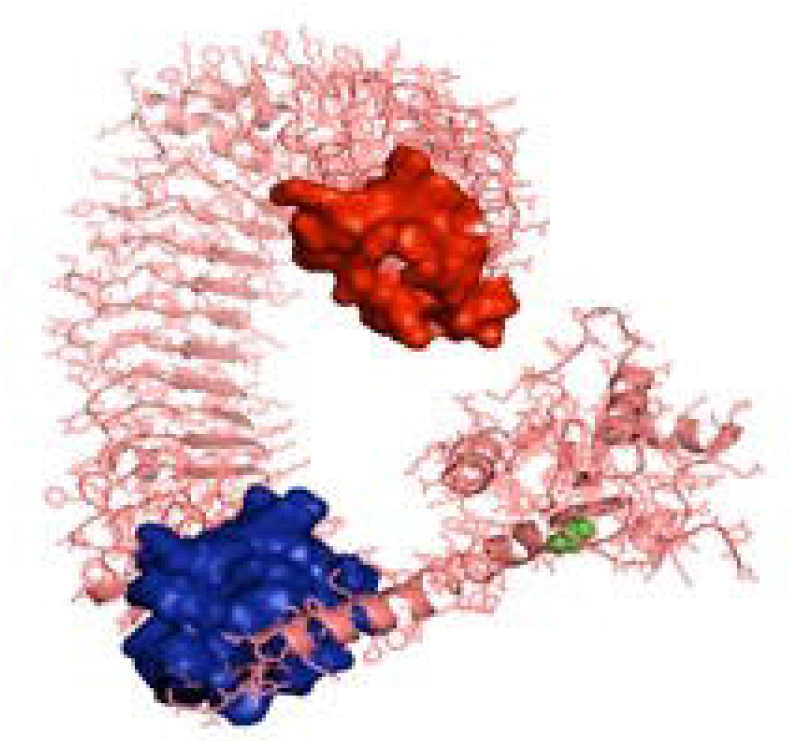
3D structure of TLR7 of duck with LRRNT, TPKRC2, GPI anchor

**Fig 3f:**
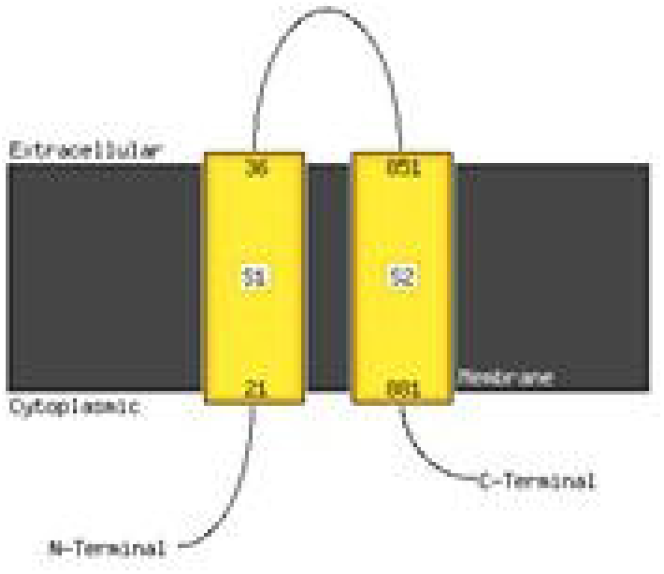
Transmembrane helix for TLR7 of duck

### Molecular docking of TLR3, RIGI and TLR7 peptide with antigenic binding site of H and N-antigen of Avian influenza virus

Binding for H and N antigen was observed for different strains of Avian influenza with RIGI, TLR7and TLR3 (Fig 4). Patchdock analysis has revealed high score for Haemagglutinin and neuraminidase antigen for H5N1 strain of Avian influenza virus. The patchdock score for H antigen for RIGI, TLR7 and TLR3 were observed to be 19920, 20532, 22880 respectively, whereas patchdock score for N antigen for RIGI, TLR7 and TLR3 were 21570, 20600, 21120 respectively as detailed in Supplementary Fig 1. The binding scores were observed to be sufficiently high. Highest score was obtained for H antigen with TLR3.

**Fig 4:**
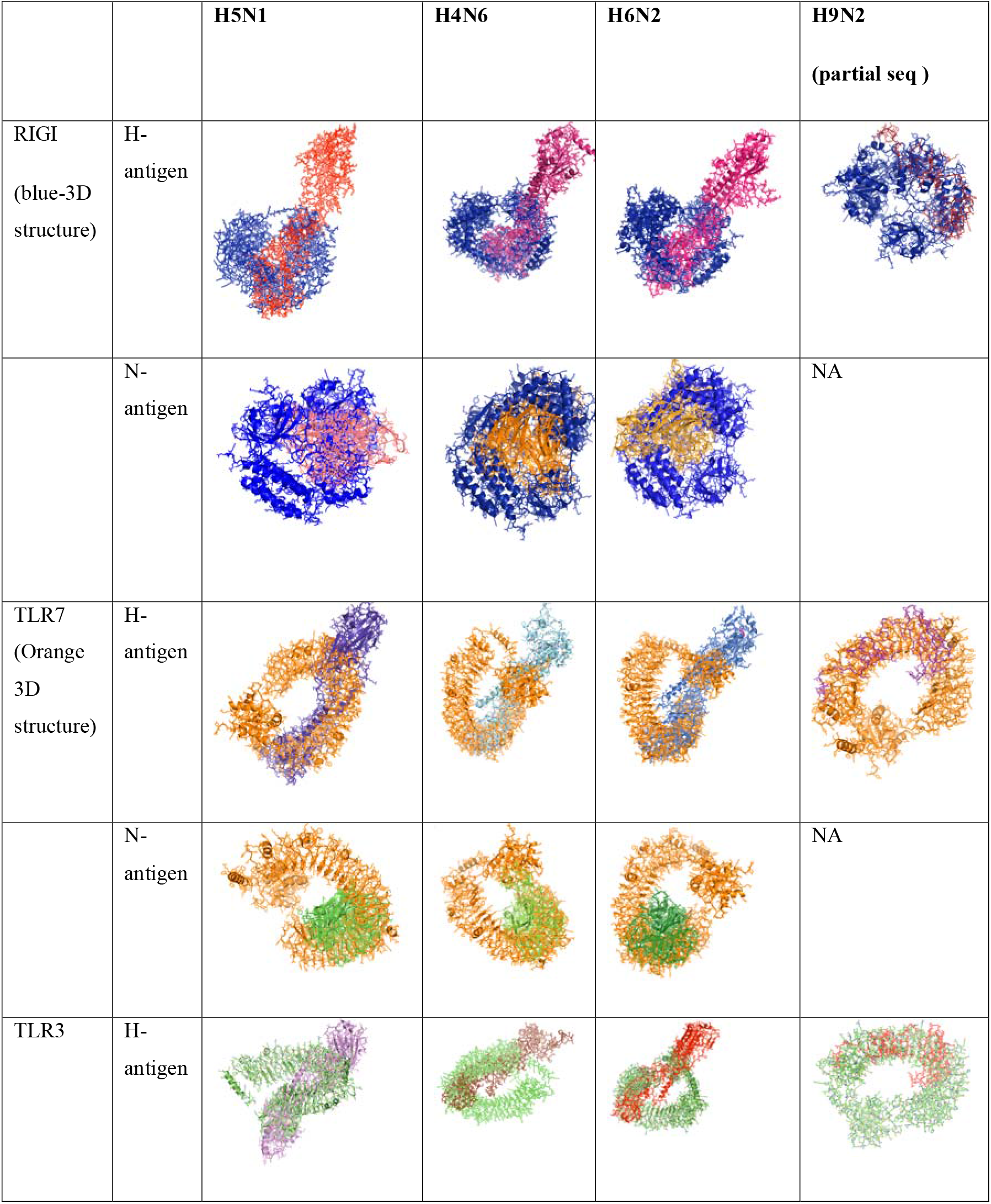
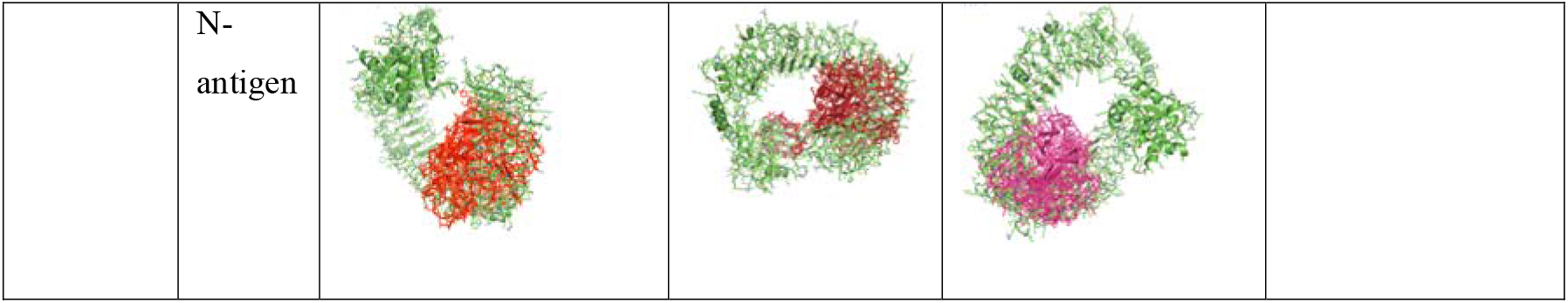
Alignment /Binding of identified immune response molecules with Haemagglutinin and neuraminidase for different strain of Avian influenza

Ligand binding is very much important for the receptor molecule. In our current study, we had studied only the surface antigens as Haemagglutinin and neuraminidase that are involved in binding with the immune molecules.

The binding of RIGI of duck with Haemagglutinin H5N1 strain of Avian influenza is being depicted with certain domains highlighted. Binding site of RIGI with H antigen of H5N1 strain of Avian influenza virus extends from 466 to 900 amino acid positions as blue spheres (Fig 5a). Red surface indicates the helicase interface domain (511-512, 515, 519 aa position of RIGI). Amino acid position 519 is a predicted site for helicase interface domain as well as site for RD interface. The site for helicase interface domain depicted as yellow surface. Other important domain within the site includes zinc binding domain depicted as green surface.

**Fig 5.** Molecular docking of RIGI with surface antigen for Avian influenza

**Fig 5a:**
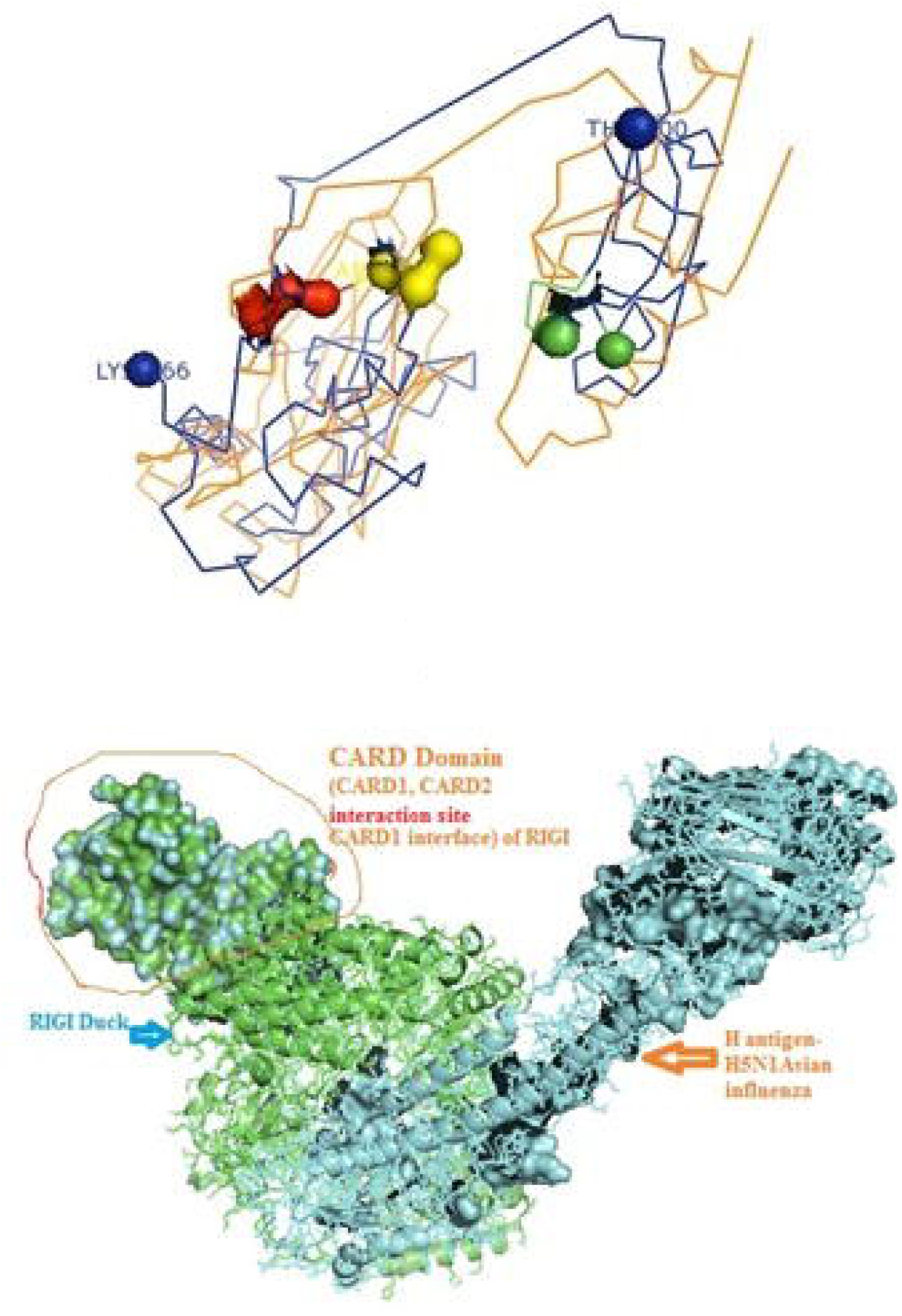
Molecular docking image RIG1 of duck with antigen (AI) ligand H antigen-binding site detection

The binding of RIGI of duck with neuraminidase H5N1 strain of Avian influenza is being depicted with certain domains highlighted. Binding site of RIGI with N antigen of H5N1 strain of Avian influenza virus extends from lysine 245 to Isoleucine 914 amino acid positions as blue spheres (Fig 5b). Fig 5b depicts only the aligned region of neuraminidase and RIGI. The site for RIG-I-C (C terminal domain of retinoic acid-inducible gene ranging from 807-921 aa position by yellow stick.

**Fig 5b:**
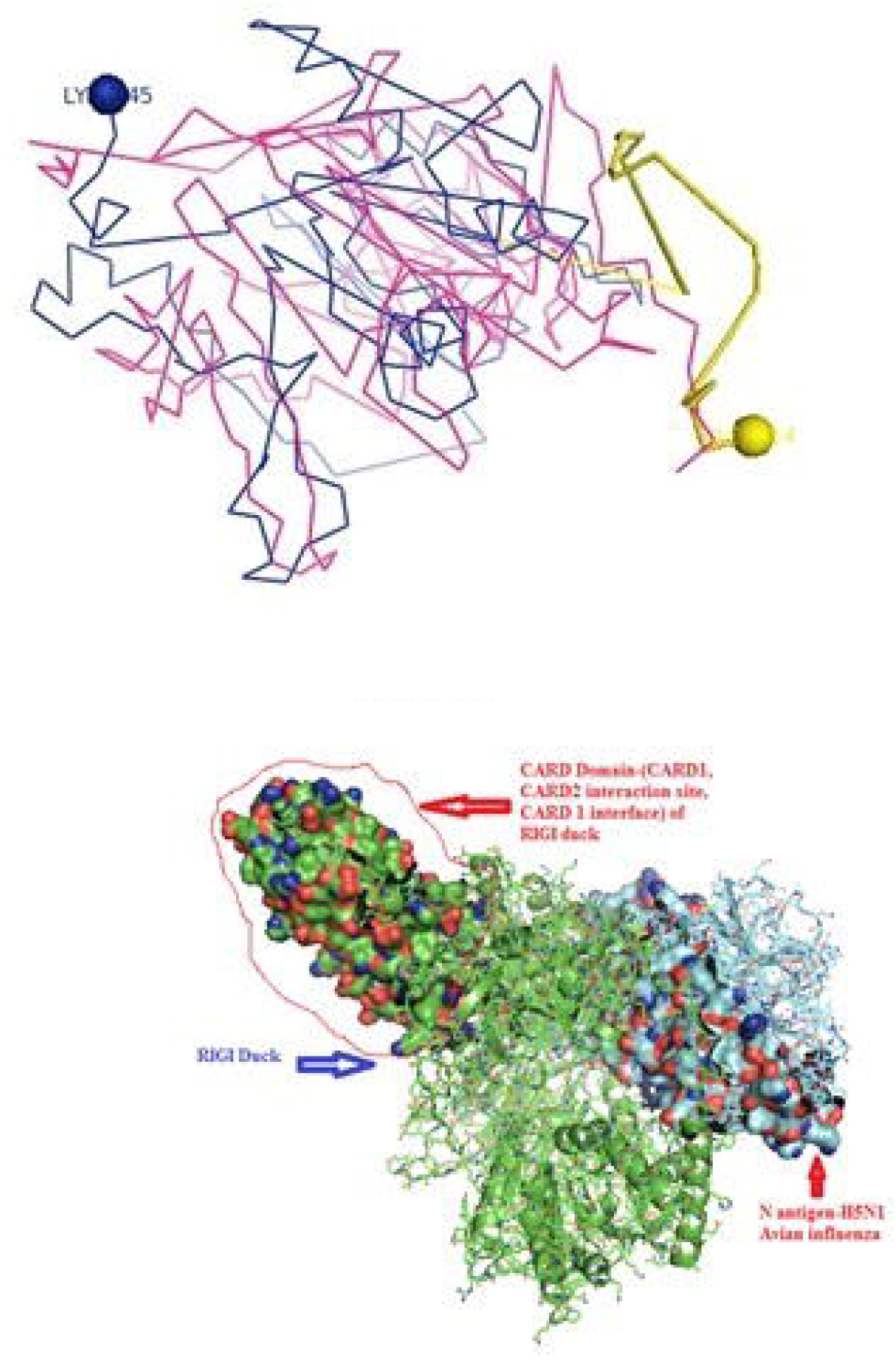
Molecular docking image RIG1 of duck with antigen (AI) NS antigen ligand(left) and binding site (right)

An interesting observation was that the CARD domains as-CARD_RIG1 (Caspase activation and recruitment domain found in RIG1), CARD2 interaction site, CARD1 interface were not involved in binding with both the surface protein Haemagglutinin and Neuraminidase of Avian influenza virus. This was proved through pdb structure of RIGI developed with Modeller software (Fig 5c and Fig 5d) respectively for H- and N-antigen.

**Fig 5c:** Duck RIGI model binding with H-antigen of Avian Influenza virus, with CARD domain-no binding for CARD with virus.

**Fig 5d:** Duck RIGI model binding with N-antigen of Avian Influenza virus, with CARD domain-no binding for CARD with virus.

The binding of TLR3 of duck with neuraminidase H5N1 strain of Avian influenza is being depicted with certain domains highlighted. Binding site of TLR3 with N antigen of H5N1 strain of Avian influenza virus extends from threonine 34 to Isoleucine 459 amino acid positions as green spheres (Fig 6a). Identifiable domains within this region includes LLR 1 to 12, site for leucine zipper, leucine-rich nuclear export signal and LRRNT. The domains within the 3D structure of TLR3 have been visualized already in Fig 1a–e.

**Fig 6a:**
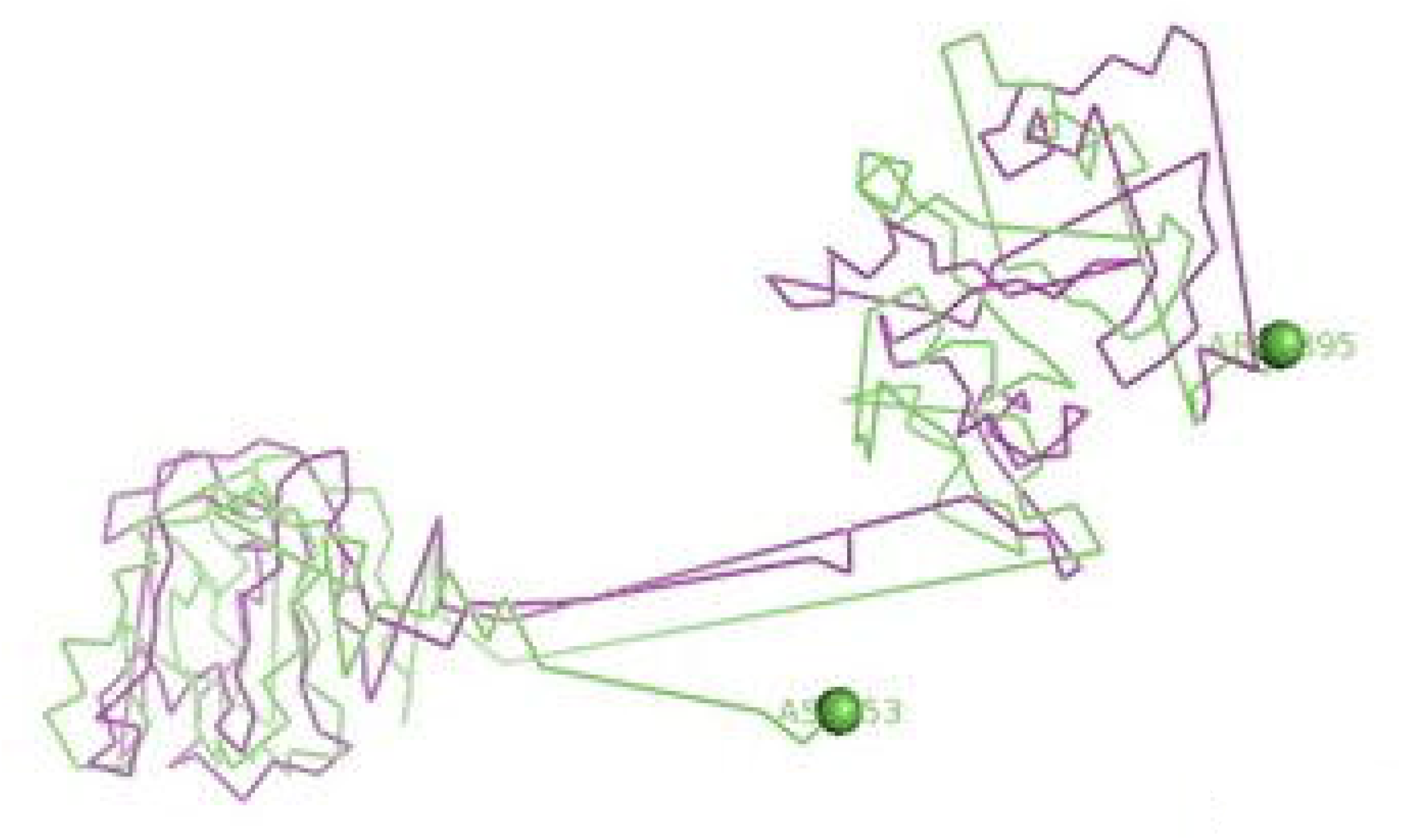
Molecular docking image RIG1 of duck with antigen (AI) ligand H antigen-binding site detection

The binding of TLR3 of duck with Haemagglutinin H5N1 strain of Avian influenza is being depicted with certain domains highlighted. Binding site of TLR3 with H antigen of H5N1 strain of Avian influenza virus extends from Asparagine 53 to Arginine 895 amino acid positions as green spheres (Fig 6b). The important domains within this region includes LRR region 1-18, site for leucine zipper, GPI anchor, leucine-rich nuclear export signal, LRRCT, site for TIR and the sites for leucine-rich receptor-like protein kinase.

**Fig 6b:**
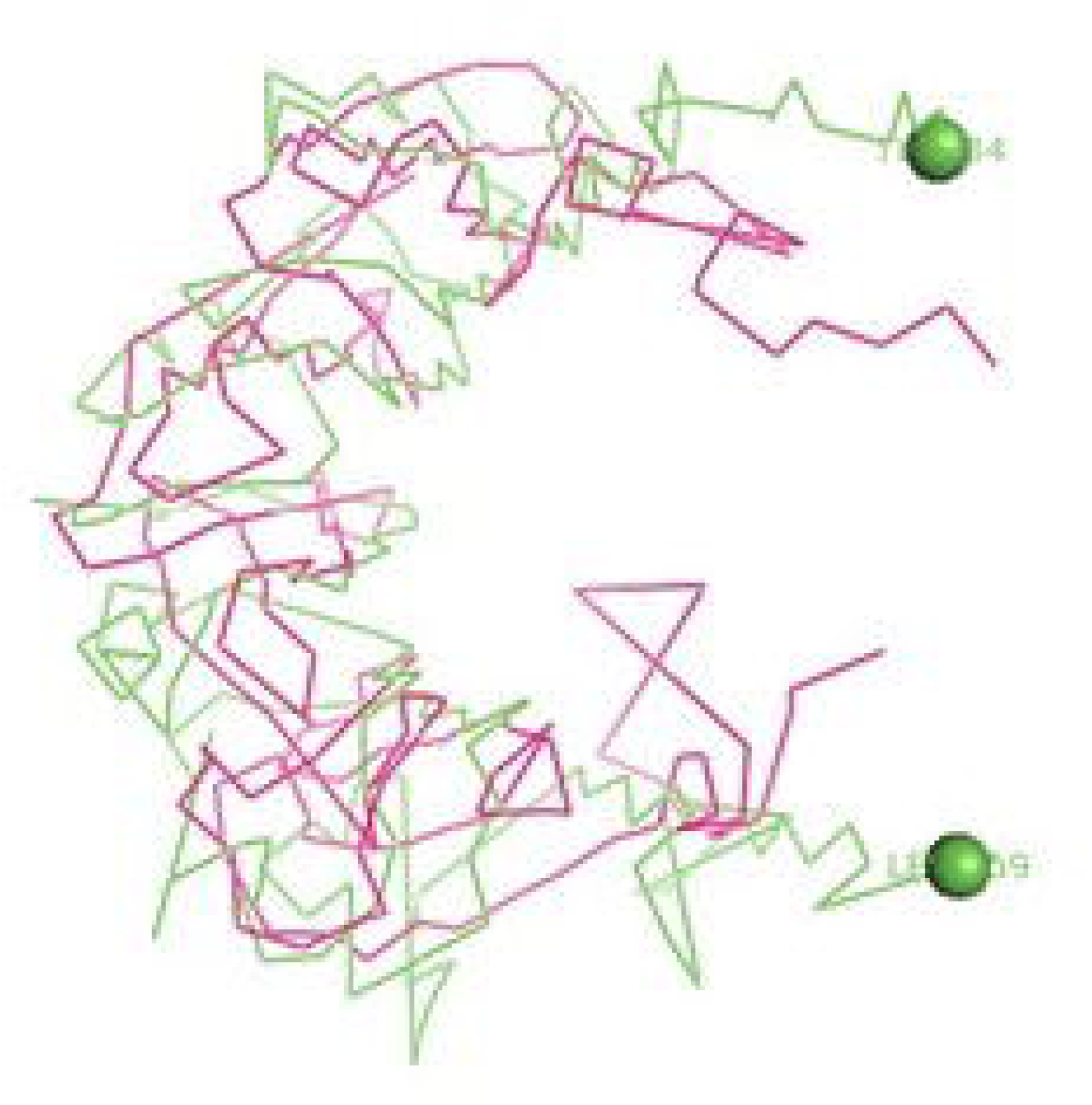
Molecular docking image RIG1 of duck with antigen (AI) NS antigen ligand(left) and binding site (right)

The binding of TLR7 of duck with neuraminidase H5N1 strain of Avian influenza is being depicted with certain domains highlighted. Binding site of TLR7 with N antigen of H5N1 strain of Avian influenza virus extends from Valine 87 to Glutamine 645 amino acid positions as orange spheres (Fig 7a). Identifiable important domains within this region includes LRR region 1-11 and domain linker sites. The detail visualization of these domain are present in Fig 3a–f in molecular visualization tool.

**Fig 7a:**
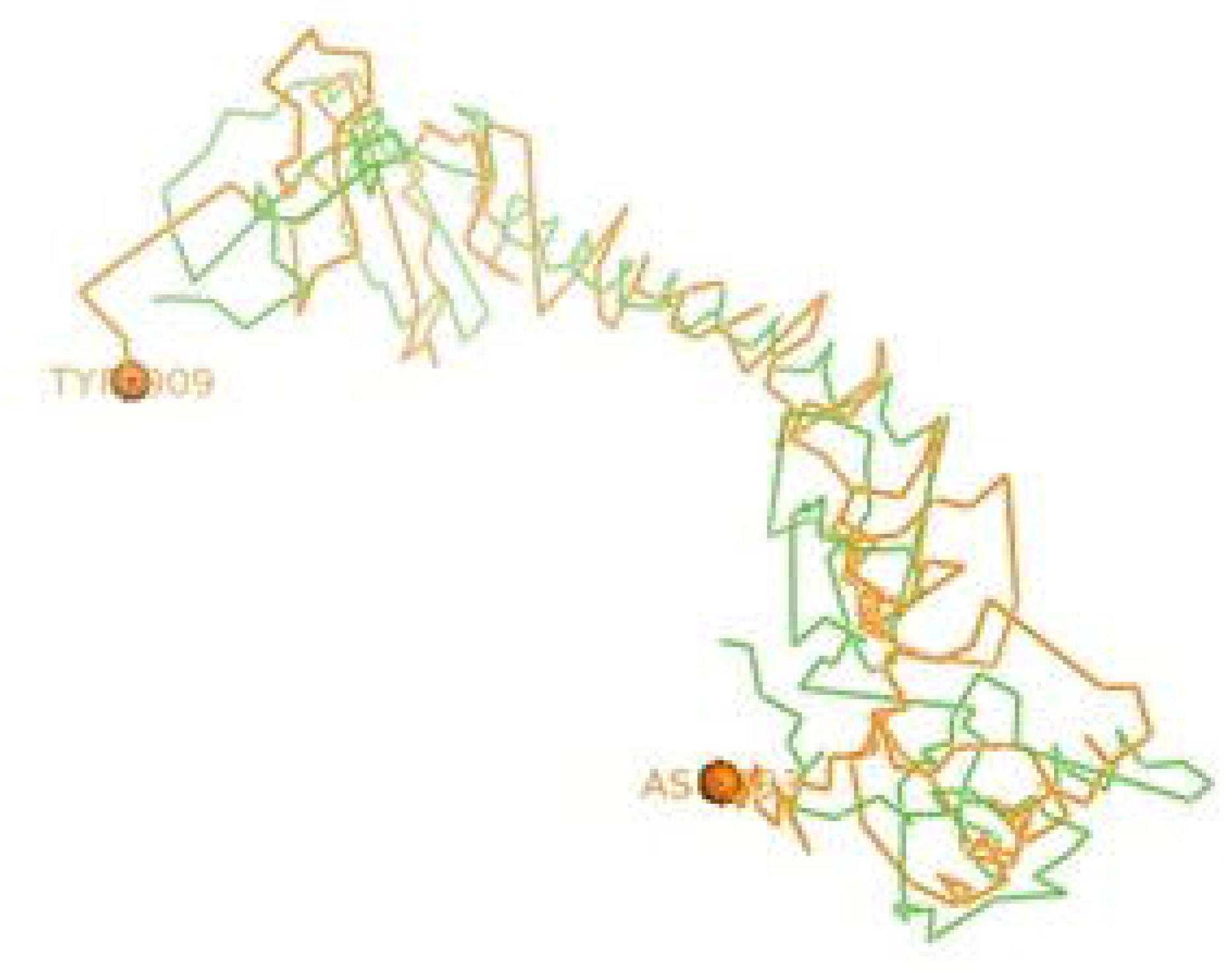
Molecular docking -tlr3 of duck with H antigen of Avian influenza

The binding of TLR7 of duck with Haemagglutinin H5N1 strain of Avian influenza is being depicted with certain domains highlighted. Binding site of TLR7 with H antigen of H5N1 strain of Avian influenza virus extends from Aspartic acid 293 to Tyrosine 909 amino acid positions as orange spheres (Fig 7b). The important domains responsible within this binding site includes LRR 7 to LRR14, domain linker sites, and cysteine-rich flanking region-C-terminal, TPKR-C2 (Tyrosine-protein kinase receptor C2 Ig like domain).

**Fig 7b:**
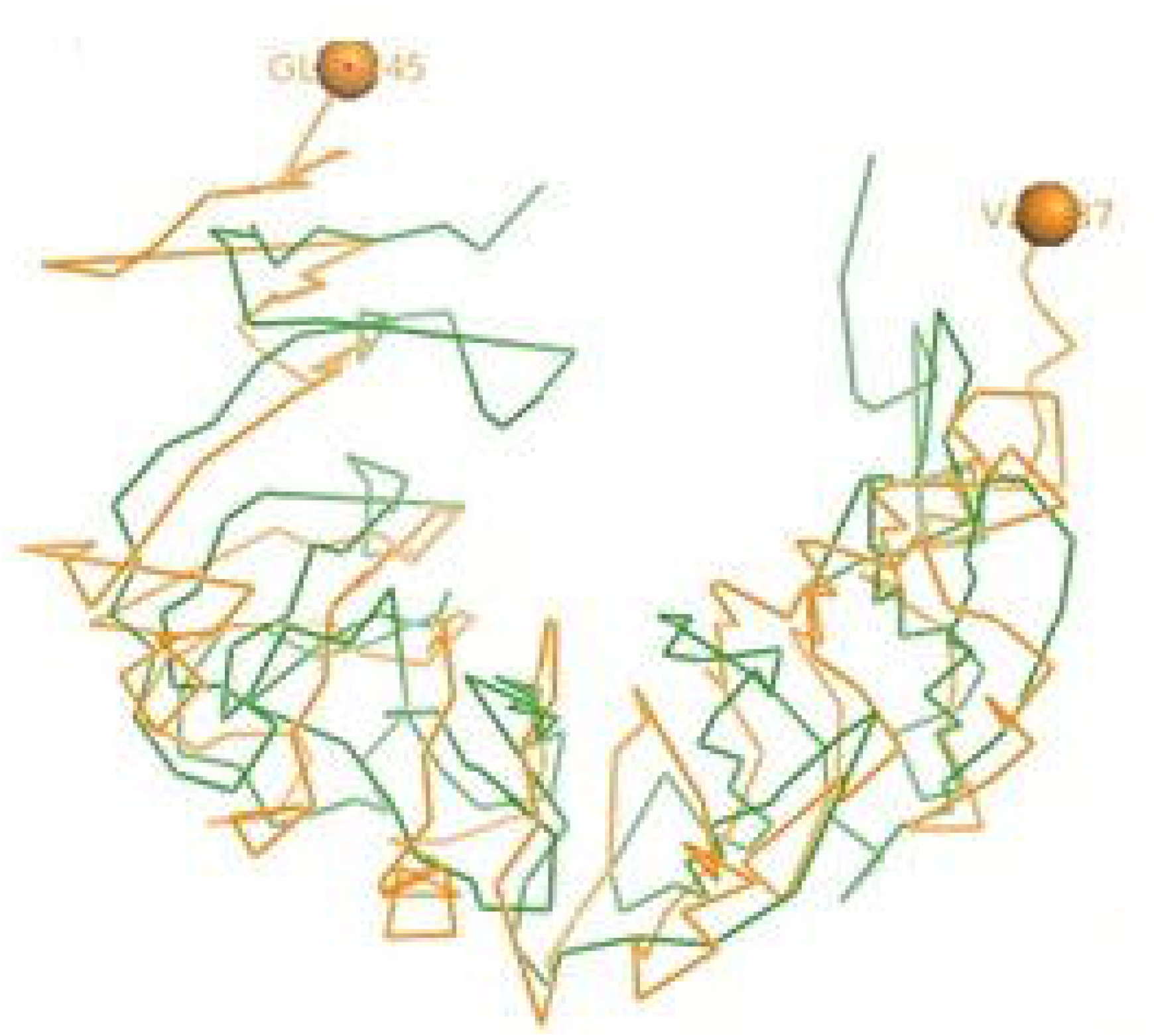
Molecular docking tlr3 of duck with NS antigen of Avian influenza

### Amino acid sequence variability and molecular phylogeny among different strains of Avian influenza

High degree of sequence variability has been observed in Fig 8a & b in Haeagglutinin segment of Avian influenza and Fig 8c and Fig 8d in neuraminidase segment of Avian influenza. H5N1 strain was observed to be clustered with H6N2 strain of Avian influenza virus. H4N6 strain comparatively less virulent causing LPAI was found to possess certain uniqueness in amino acid sequence. Deletions of four consecutive amino acids at positions 65-68, 140-141 were observed in Haemagglutinin. Likewise insertion mutations were also observed at amino acid positions 19-24, 77-79 in Haemagglutinin. Cysteine residues were observed to be conserved across the strains (Fig 8b).

**Fig 8a:**
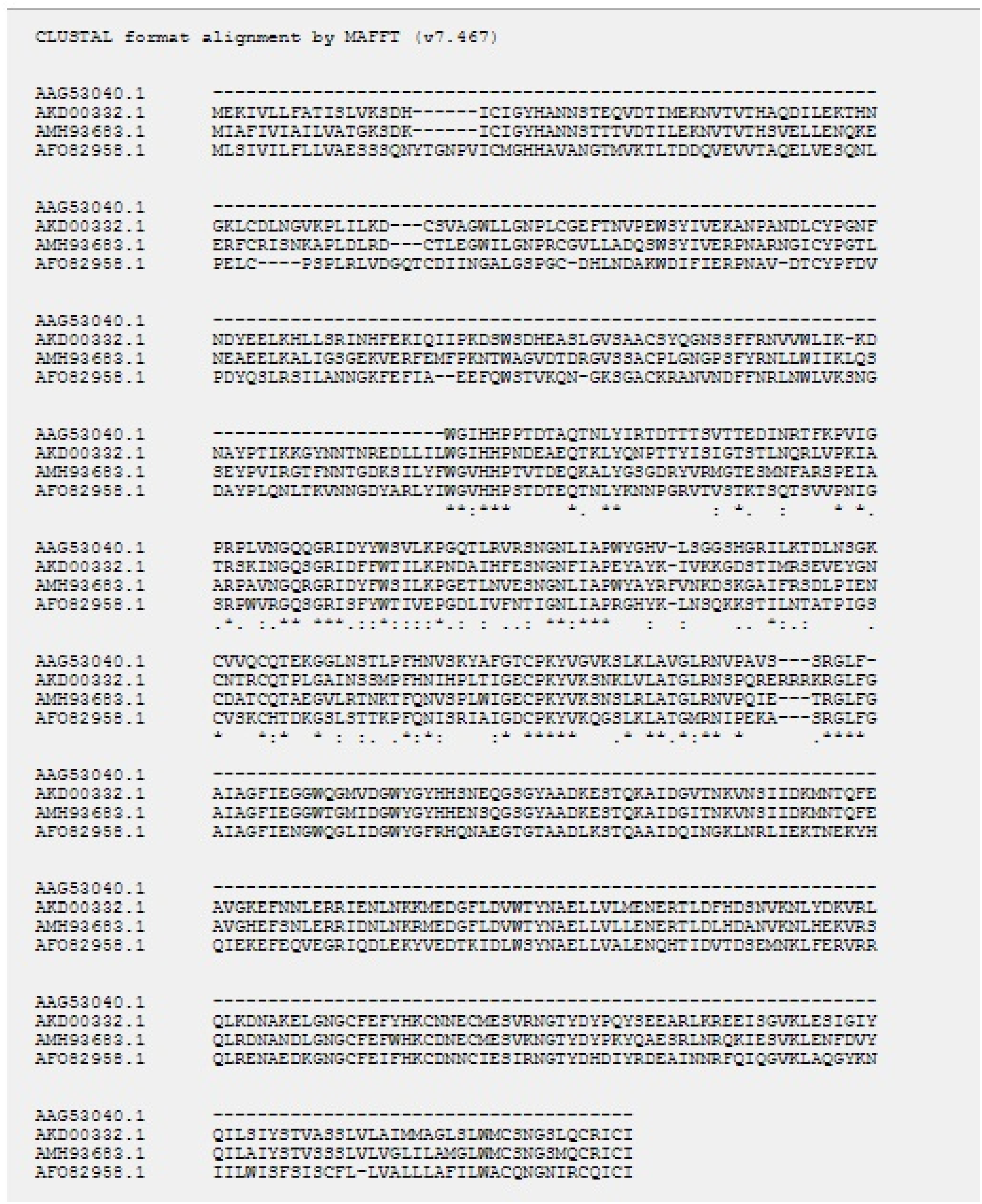
Molecular docking TLR7 of duck (red) with Avian influenza(blue) H antigen-binding site

**Fig 8b:**
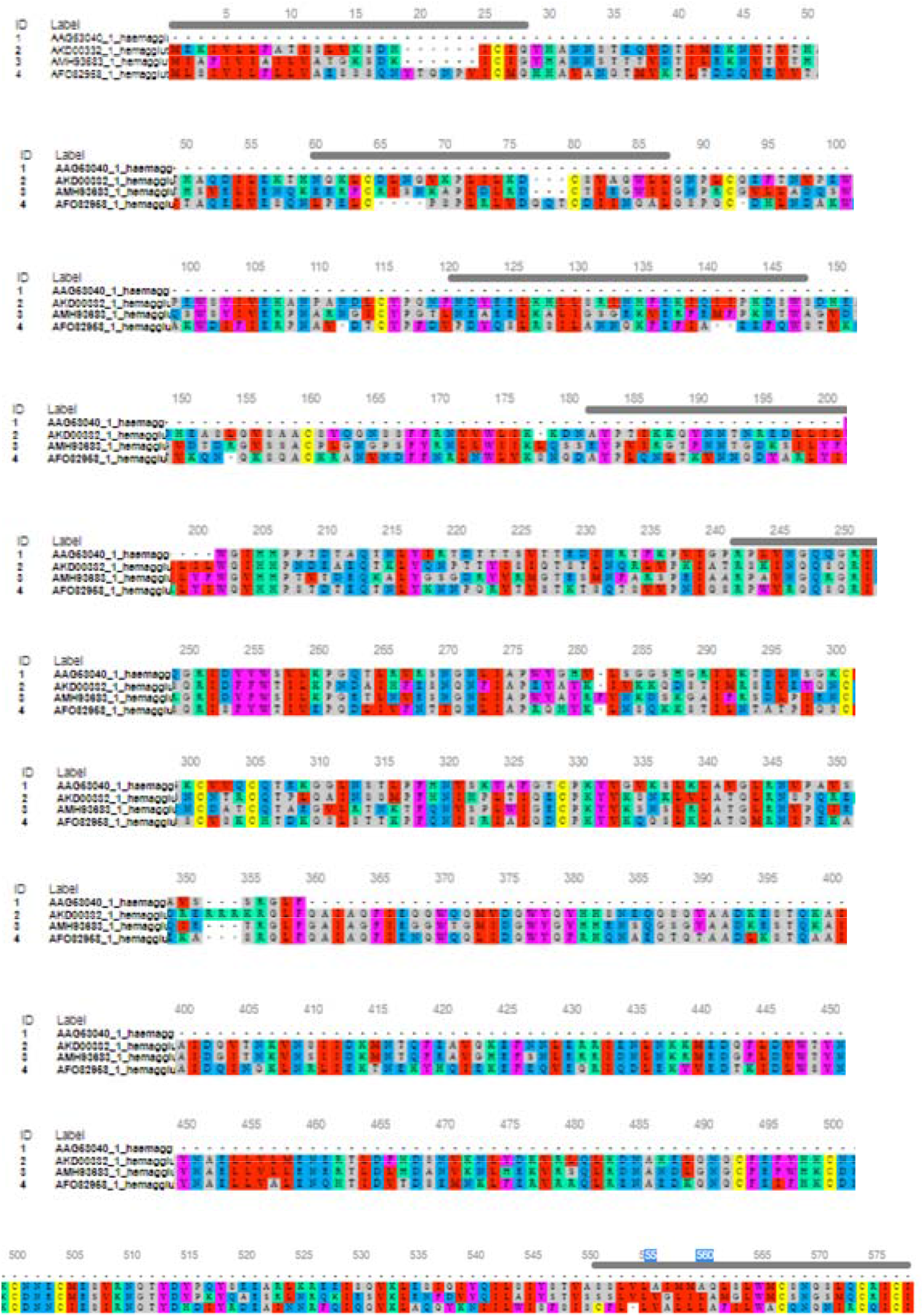
PatchDock TLR7 of duck (red)/ avian influenza(blue) NA antigen

**Fig 8c.**
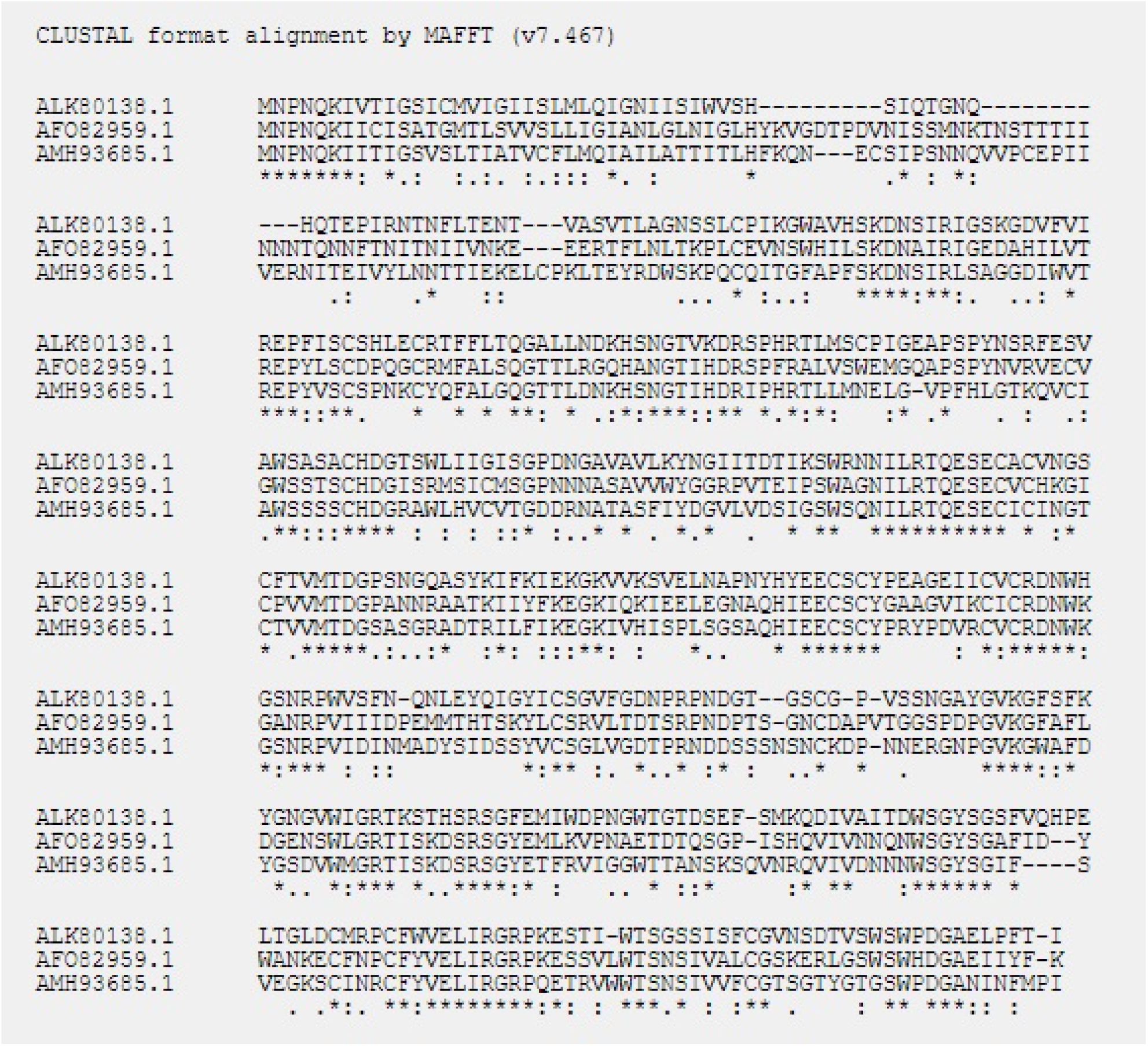

**Fig 8d.**
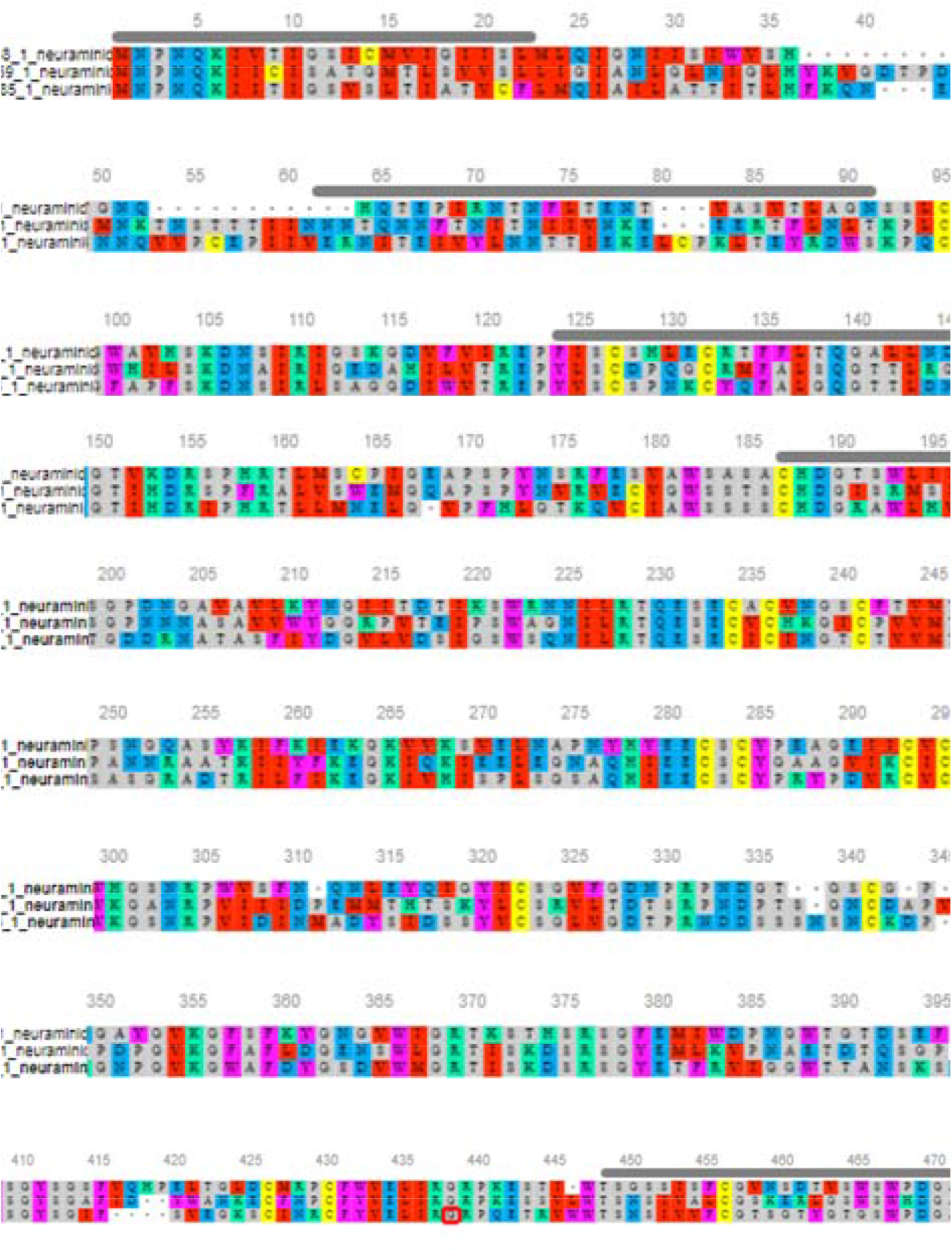

However in case highly pathogenic H5N1 depicts certain deletion mutations at amino acid positions 37-45, 53-63, 80-82 of neuraminidase (Fig 8d).

### Comparative structural analysis of TLR3 and TLR7 of duck with respect to chicken

Ducks were reported to be genetically more resistant to chicken, particularly in terms of viral infections. Accordingly, the structural alignment of 3D structure of TLR3 of duck with chicken has been described (Fig 9a). 3D structural alignment of TLR7 of duck with chicken have been visualized (Fig 9b). It was not possible to study the structural alignment of Duck RIG-I, since RIGI was not expressed in chicken.

**Fig 9A.**
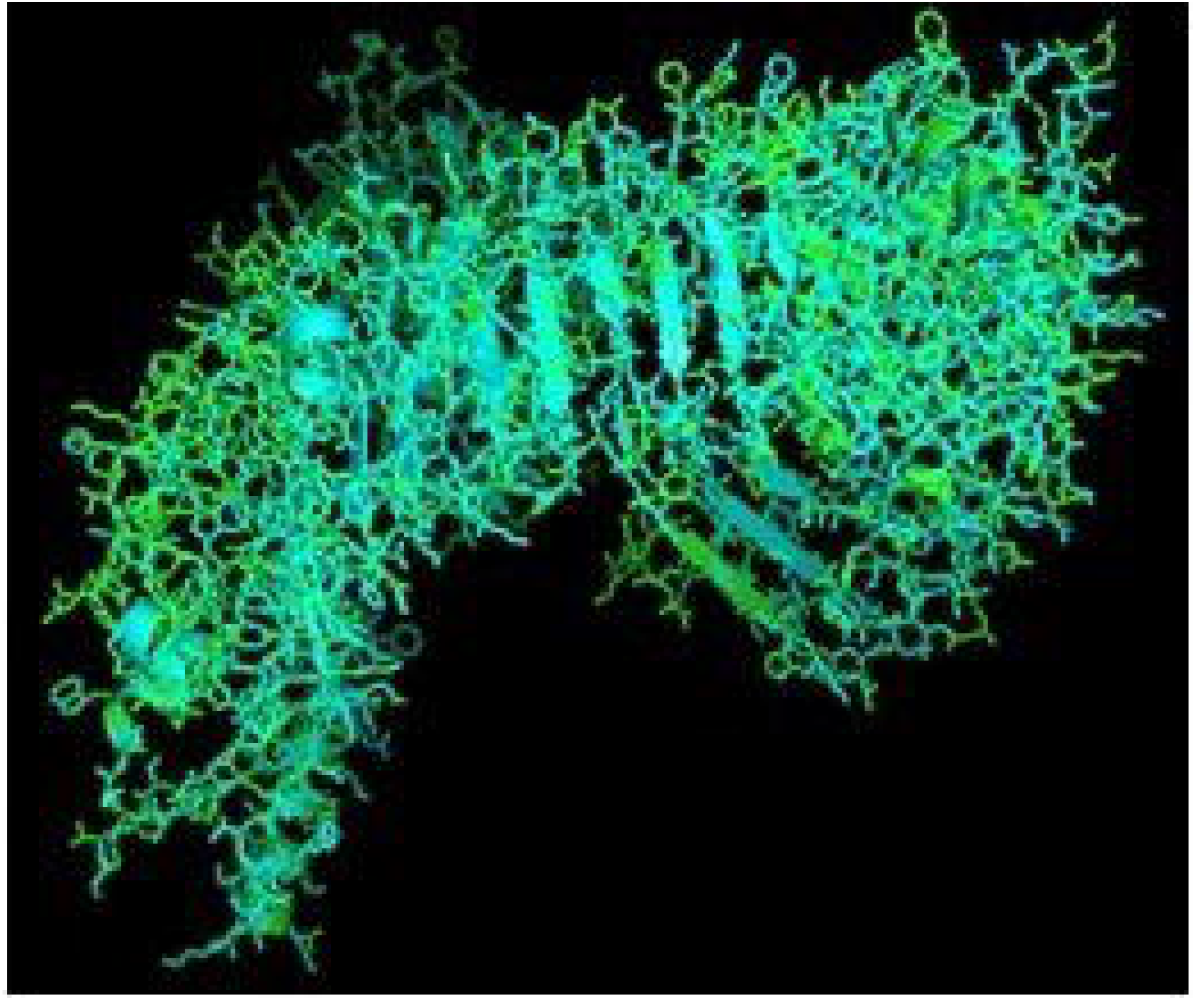
Alignment report for Haemagglutinin antigen for different strains for Avian influenza

**Fig 9B.**
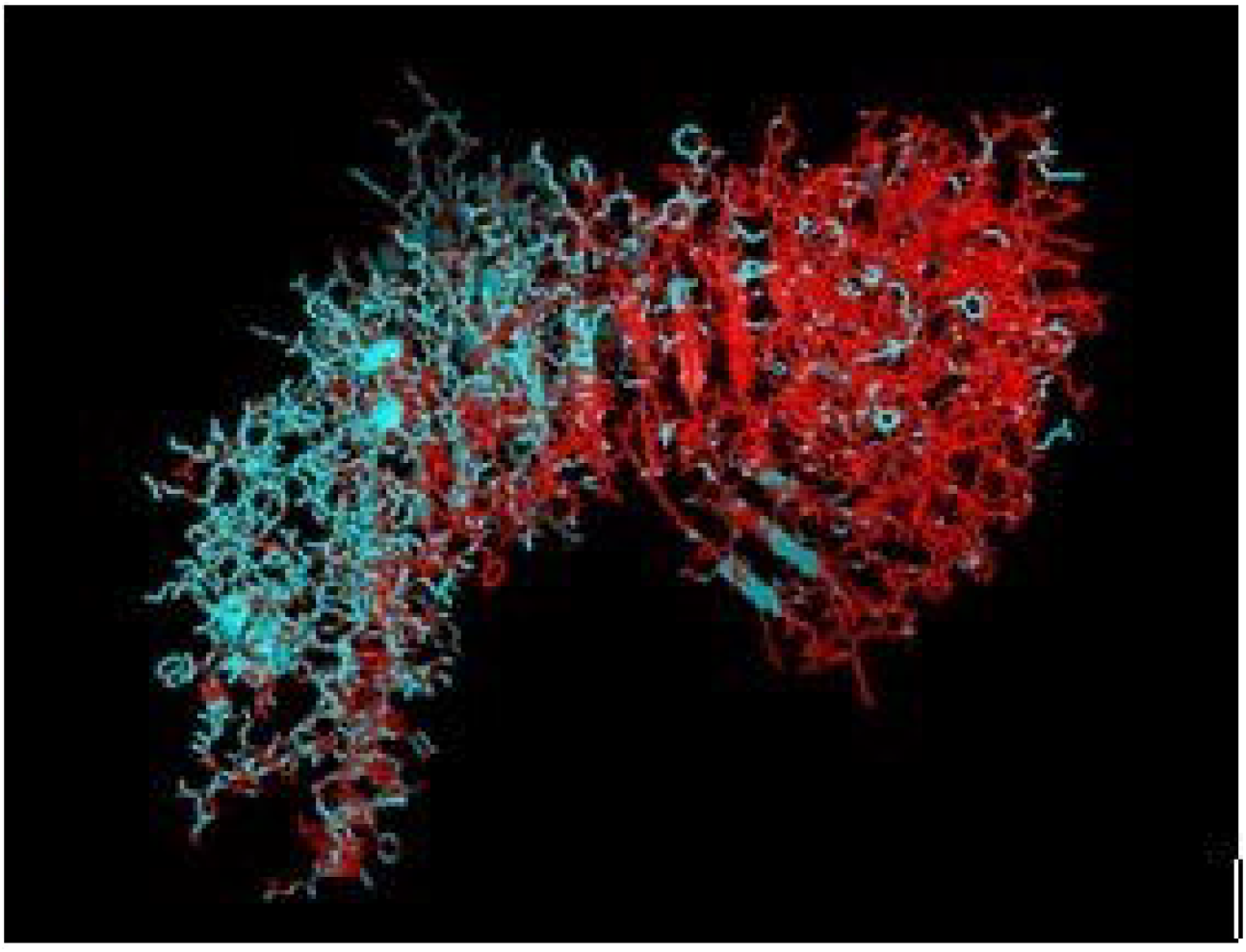
Alignment report for Haemagglutinin antigen for different strains for Avian influenza by MAFFT software.

**Fig 9C.** Alignment report for neuraminidase antigen for different strains for Avian influenza

**Fig 9D.** Alignment report for neuraminidase antigen for different strains for Avian influenza by MAFFT software.

**Fig 9E.** Phylogenetic analysis for different strains for Avian influenza w.r.t. Haemagglutinin antigen.

The sites for non-synonymous mutations have been depicted for TLR3 gene in duck with respect to other poultry species as chicken, goose and guineafowl (Table 2). 51 sites for amino acid substitutions have been detected ranging from amino acid position 42 to 766 in duck with respect to other poultry species, which actually contribute to changes in functional domains of TLR3. Comparison of TLR3 of duck with chicken actually revealed 46 sites of amino acid substitution resulting due to non-synonymous mutations, which are of much importance to our present study. Most of the substitutions caused changes in Leucine rich repeats, which is an inherent characteristic for Pattern recognition receptor as TLR2. 20 sites of amino acid substitutions were identified that were specific for Anseroides (Duck and goose).

### Protein-protein interaction network depiction for TLR3 and TLR7 with respect to other functional proteins

Interaction of TLR3 with other proteins has been depicted in fig 10a with STRING analysis. Interaction of TLR7 with other protein of functional interest has been depicted in Fig 10b. KEGG analysis depicts a mode of the defense mechanism of influenza A and the possible role of antiviral molecules in combating the infection and the role of antiviral molecules through TLR signaling pathway.

**Fig 10.**
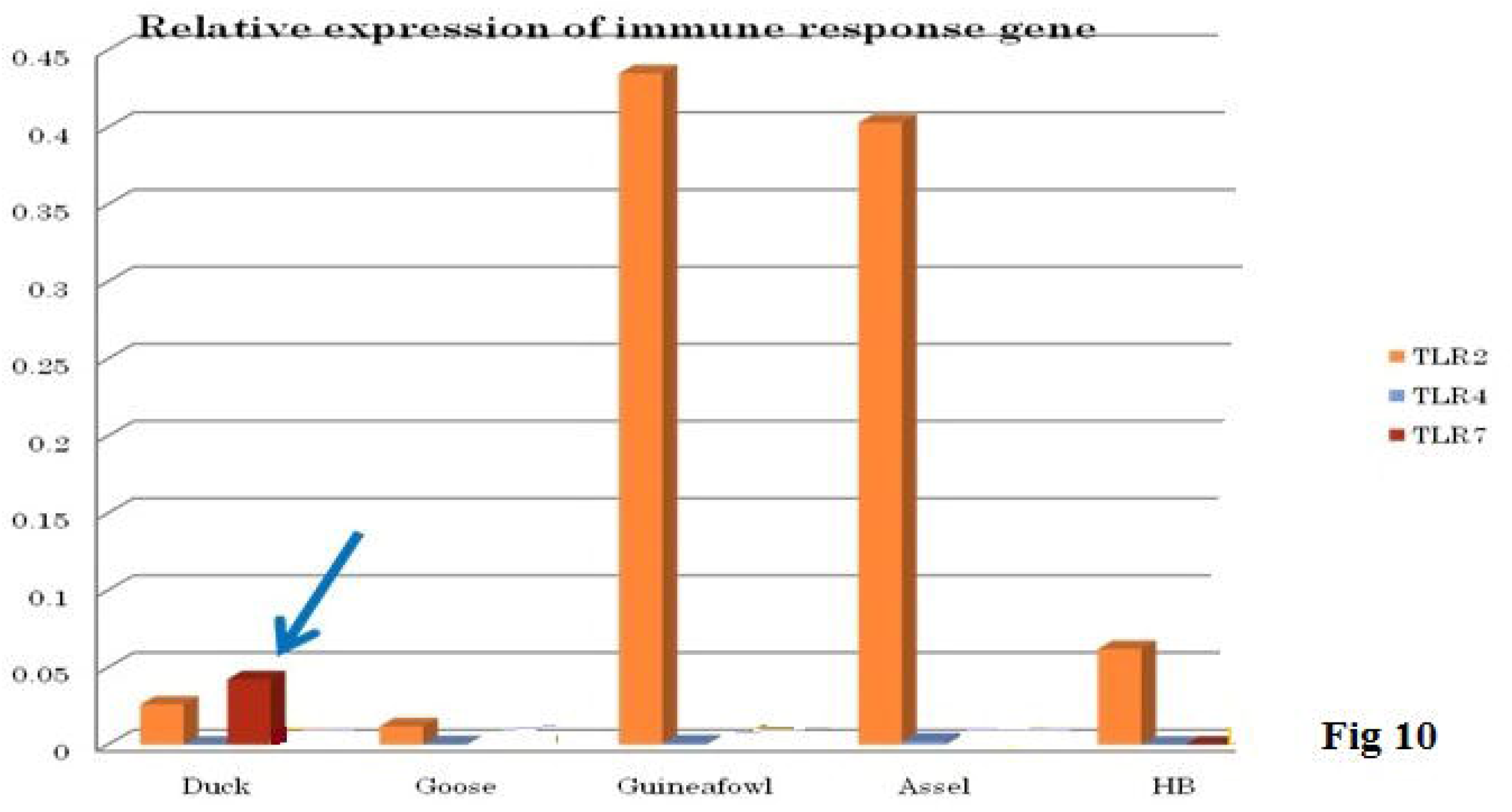

**Fig 10a:**
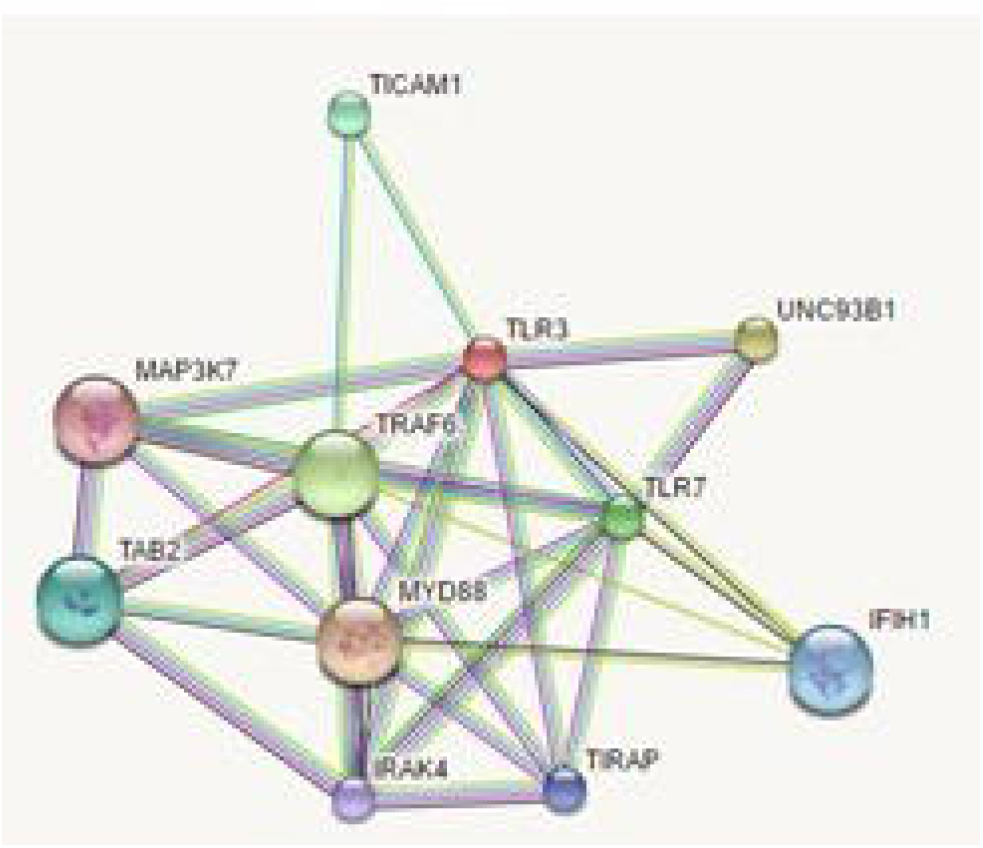
Comparison of TLR3 of duck and chicken

**Fig 10b:**
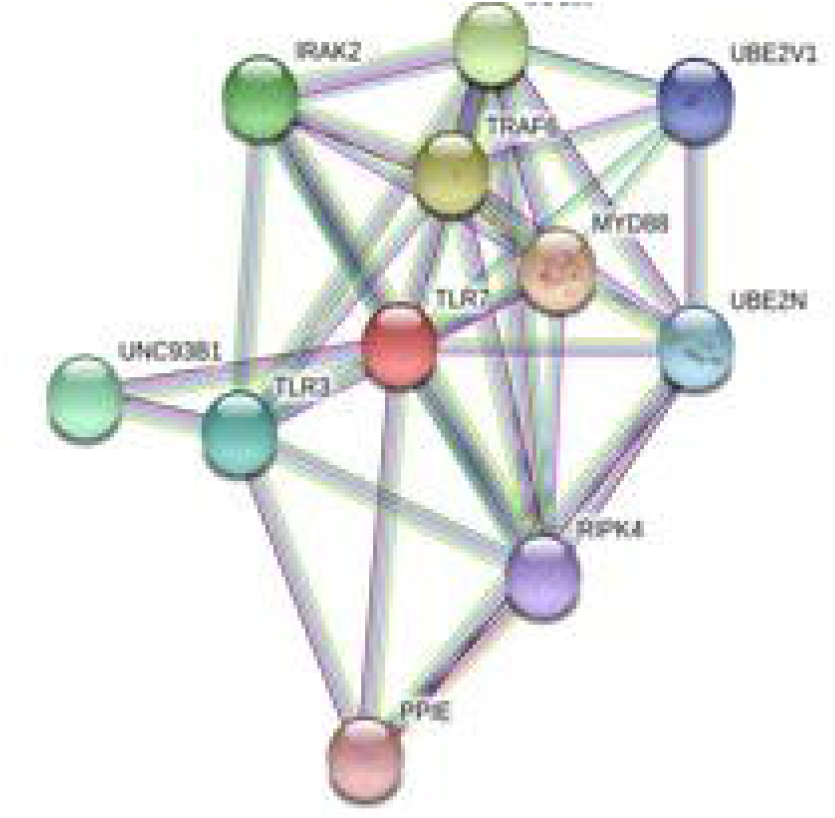
Comparison of TLR7 of duck and chicken

### Differential mRNA expression pattern of TLR7 and other TLR gene of duck with respect to chicken and other poultry species

We conducted differential mRNA expression profiling of TLR2, TLR4, TLR7. TLR2 and TLR4 expression profiling was observed to be better in indigenous chicken (Aseel and Haringhata Black) and guineafowl in comparison to Anseroides (Duck and Guineafowl). Both TLR2 and TLR4 are known to impart antibacterial immunity. Quantitative mRNA expression analysis clearly depicts TLR7 gene expression was definitely better in duck compared to other poultry species as Goose, guineafowl and indigenous chicken breed (Aseel and Haringhata Black chicken) (Fig 11). This gives an indication that better immune response of indigenous duck may be due to increased expression level of TLR7, which confer antiviral resistance (Fig 11).

**Fig 11a:** String analysis revealing molecular interaction of TLR3

**Fig 11b:** String analysis revealing molecular interaction of TLR7

### Phylogenetic analysis of indigenous ducks with other poultry species and other duck population globally

With an aim for the identification of the status of molecular evolution of duck, the indigenous duck gene sequence of West Bengal, India was compared with other duck sequences globally. Phylogenetic analysis was analysis with respect to TLR7 (Fig 12a) and TLR3 (Fig 12b). Phylogenetic analysis revealed that ducks of West Bengal was observed to be genetically more closely related to the duck population of china (Fig 12a). Ducks were observed to be genetically closest to goose (Fig 12b). Chicken, quail, and Turkey were observed to be genetically distinct from duck (Fig 12b).

**Fig 12:**
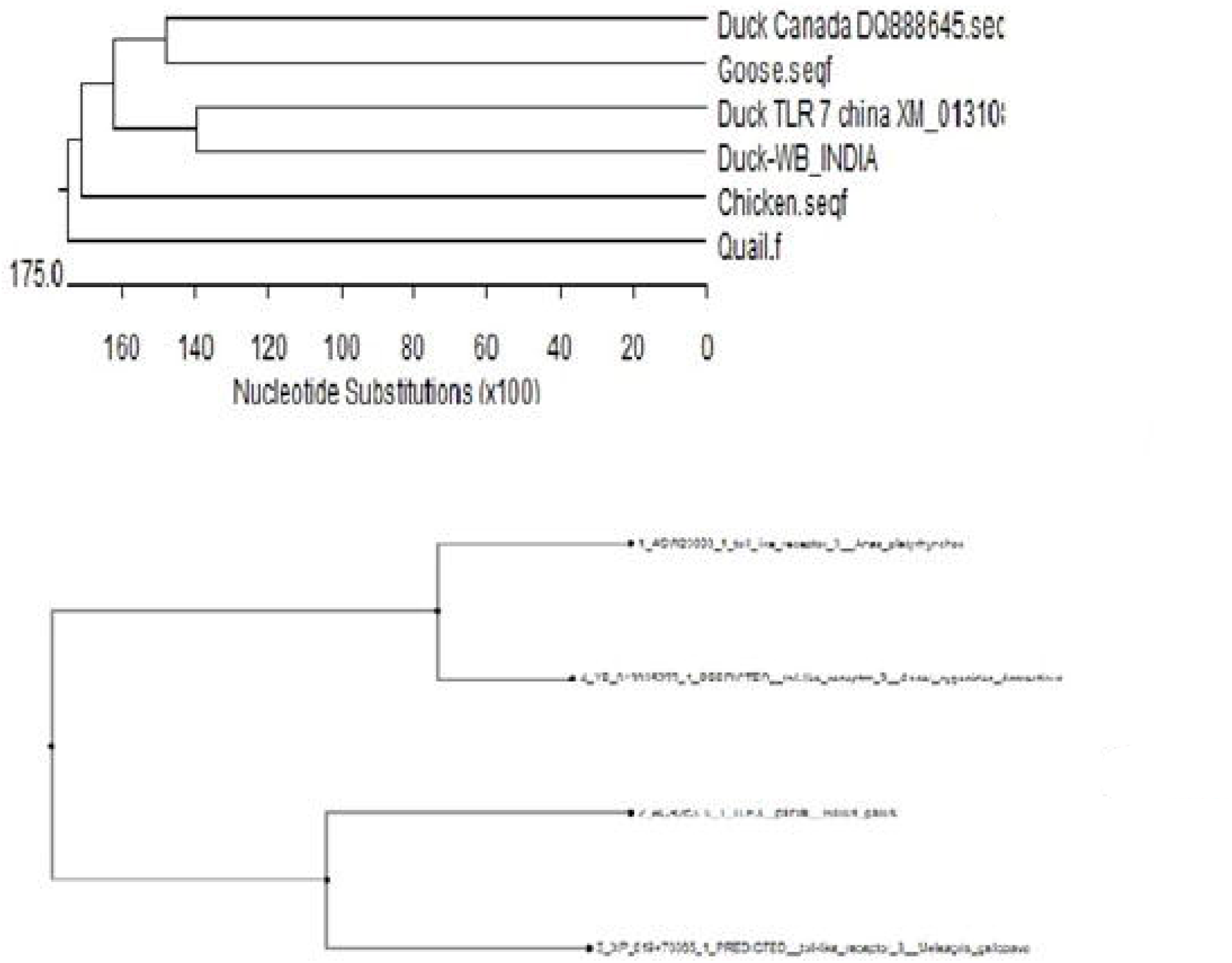
KEGG analysis for defense mechanism against Influenza A infection

**Fig 13:** TLR signalling pathway (KEGG) (TLR3 and TLR7)

**Fig 14:** Differential mRNA expression profile of immune response genes of duck with other poultry species

**Fig 15A:** Molecular evolution of ducks reared globally in relation to other poultry species (Based on TLR7)

**Fig 15B:** Molecular evolution of ducks reared globally (Based on TLR3)

## Discussion

Indigenous duck population were characterized to be very hardy, usually asymptomatic to common avian diseases. But there is a paucity of information regarding the systemic genetic studies on duck involved in its unique immune status. It is evident that duck possesses some unique genetic makeup which enables it to provide innate immunity against viral infection, particularly avian influenza.

In this current study, we identified three immune response molecules, earlier known to have immunity against viral infection as RIGI, TLR7, and TLR3. We characterized these proteins of duck, attempted to identify the SNPs or variations in nucleotide among duck and chicken. Through molecular docking, the most promising IR molecules conferring innate immunity against avian influenza have been identified, later on confirmed through wet lab study as differential mRNA expression profiling. We attempt to explore the unique genetic constitution of duck immune response with respect to that of chicken. However RIGI was reported to be expressed only in duck, not in chicken, hence comparison was not available. TLR7 expression profile was observed to be significantly better in duck in comparison to chicken and other poultry species, indicative of better antiviral immunity in ducks. 53 non-synonymous mutations with amino acid variations were observed while comparing amino acid sequence of duck with other poultry species, including chicken, most of which are confined to LRR domain. We had already depicted that LRR is an important domain for pathogen binding site as in this case of Avian influenza. For the effective antiviral activity, binding of viral protein with the immune response molecule is the primary criteria. Leucine rich repeats (LRRs) were observed to be important domain involved in binding with Haemagglutinin and neuraminidase surface protein in case of both TLR3 and TLR7. Similar studies have also reported LRR as important domain against bacterial infections in case of CD14 molecule in cattle^**19**^, goat^**17, 18**^and buffalo ^**20,21**^. Other important domains identified were LRRNT, LRRCT, site for TIR and the sites for leucine-rich receptor-like protein kinase, including certain post translational modification sites. Similar reports were also identified in different species ^**17,18,19,20,21**^. An important observation identified was that although CARD domain was believed to be an important binding site for RIGI for some identified virus, it has no binding ability with Avian influenza virus. Other studies have reported the role of CARD domain in binding with MAVS domain as a part of antiviral immunity^**42,43**^.

TLR3 (CD283 or cluster of differentiation 283) is a pattern recognition receptor rich in Leucine-rich repeats as revealed in duck TLR3 of the current study. Other important domains include LRRNT, LRRCT, TIR, leucine-rich receptor-like protein kinase, leucine zipper, GPI anchor, leucine-rich nuclear export signal. The other sites for post-translational modification as observed were N-linked glycosylation, casein kinase, phosphorylation, myristylation, phosphokinase phosphorylation. Variability of amino acids in important domains was observed for duck TLR3 compared to that of Chicken, Goose, and Turkey. It was observed that genetic similarity between duck and goose was more compared to that of chicken and Turkey. Some amino acids have been identified which are conserved for ducks and denote for important domains as LRR, LRRCT, TIR. In the LRRCT domain, valine is present in a duck in contrast to alanine in chicken, turkey, and goose. The current study identified 45 sites of non-synonymous substitutions between duck and chicken, which affects important domains for TLR3 as pattern recognition receptor. Although it is the first report of characterization of TLR3 in indigenous duck, earlier studies were conducted in Muscovy duck, when full-length cDNA of TLR3 was characterized to be of 2836 bp encoding polypeptide of 895 amino acids^44^. The characterization of the deduced amino acid sequence contained 4 main structural domains: a signal peptide, an extracellular leucine-rich repeats domain, a transmembrane domain, and a Toll/IL-1 receptor domain^44^, which is in agreement to our current study. It is to be kindly noted that TLR3 is a PRR, with the secondary structure being visualized as helix, loop, and sheet with the sites for disulfide bond being depicted in blue spheres. It recognizes dsRNA associated with a viral infection, and induces the activation of IRF3, unlike all other Toll-like receptors which activate NF-κB. IRF3 ultimately induces the production of type I interferons, which aids in host antiviral immunity^45^. In the current study we observed sites for leucine zipper in duck TLR3, which is an inherent characteristics for dimerization. Earliers studies have also reported that TLR3 forms a large horseshoe shape that contacts with a neighboring horseshoe, forming a "dimer" of two horseshoes^**46**^. As already explained that glycosylation is an important PTM (post-translational modification site), it acts a glycoprotein. But in the proposed interface between the two horseshoe structure, two distinct patches were observed rich in positively charged amino acids, may be responsible for the binding of negatively charged viral dsRNA.

RIG1 (also known as DEAD-box protein 58 (DDX58) is an important molecule conferring antiviral immunity. Various important domains have been identified as CARD_RIG1 (Caspase activation and recruitment domain found in RIG1), CARD2 interaction site, CARD1 interface, helicase insert domain, double-stranded RNA binding site, RIG-I-C (C terminal domain of retinoic acid-inducible gene, RIG-I protein, a cytoplasmic viral RNA receptor), RD interface, zinc-binding domain, RNA binding. CARD proteins were observed to be responsible for the recognition of intracellular double-stranded RNA, a common constituent of a number of viral genomes. Unlike NLRs, these proteins, RIG-I contain twin N-terminal CARD domains and C-terminal RNA helicase domains that directly interact with and process the double-stranded viral RNA. CARD domains act through the interaction with the CARD motif (IPS-1/MAVS/VISA/Cardiff) which is a downstream adapter anchored in the mitochondria^**47,48**^.

Through *in silico* alignment study, it was clearly observed that TLR3 binds with H antigen of the avian influenza virus (H5N1). It was known that avian influenza virus (H5N1) strain contain 100-200nm spherical, enveloped, 500 projecting spikes containing 80% haemagglutinin and 20% neuraminidase^**49**^, with genome being segmented antisense ssRNA. In order to combat the infection, TLR3 binds with the haemagglutinin spikes of the influenza virus.TLR3 gene was observed to have a role to combat against Marek’s disease^**50**^. Seven amino acid polymorphism sites in ChTLR3 with 6 outer part sites and 1 inner part site ^**51**^. TLR3 cannot act alone. It acts while interacting with a series of molecules as TICAM1, MAP3K7, TAB2, TRAF6, Myd88, IRAK4, IFIH1, and even TLR7. TLR3 acts through RIG1 like receptor signaling pathway and acts through TRIF^**50**^.

RIG1 is an important molecule, which is only expressed in ducks, not in a chicken. Duck RIGI transfected cells were observed to recognize RIG-I ligand and a series of antiviral genes were expressed as IFN-β, MX1, PKR, IFIt5, OASI and consequently HPAIV (Highly pathogenic avian influenza virus) titers were reduced significantly^**52, 53**^. RIG-I belongs to the IFN-stimulated gene family and it acts through RIG-I like receptor signaling pathway. RIG1 detects dsRNA virus in the cytoplasm and initiates an antiviral response by producing Type-I and Type III IFN, through the activation of the downstream signaling cascade. RIG-I is an IFN-inducible viral sensor and is critical for amplifying antiviral response^**54,55**^. Although RIG1 expression is absent in chicken, it can produce INFα by another pathway. It has been observed that IFN-β expression upon influenza infection is mediated principally by RIG1^**56**^. INFα expression induced by chicken is unable to protect the host from avian influenza infection as IFN-β, produced in ducks ^**57**^. This may be one of the major reason why ducks are resistant to HPAIV, but not chicken. Since RIG1 gene is not expressed in chicken, a comparative study was not possible. Avian influenza virus was observed to have surface glycoproteins as haemagglutinin and neuraminidase spikes on its outer surface^**58**^. It was observed that duck RIG1 can bind with both H antigen and NA antigen of Avian influenza virus. It is interesting to note how RIGI acts on virus and causes destruction. RIG-I acts through RIG-I like signaling pathway, secretes NRLX1, IPS1, leading to the production of IKKβ, which in turn causes secretion of NFκβ, Iκβ^**59**^. These substances ultimately cause viral myocarditis and the destruction of the virus^**60**^. A sequence of reactions occurs as high fever, acute respiratory distress syndrome, chemoattraction of monocytes and macrophages, T-cell activation and antibody response^**61**^.

TLR7 is another important molecule responsible for antiviral immunity, it recognizes single-stranded RNA as the genetic material. It acts through the Toll-like receptor signaling pathway. However, from the current study, it was observed that TLR7 bind well with NA antigen. The identified domains for TLR7 are mainly LRR (Leucine-rich repeat), TIR, cysteine-rich flanking region, LRRNT (Leucine-rich repeat N-terminal domain), TPKR-C2 (Tyrosine-protein kinase receptor C2 Ig like domain). TLR7 releases Myd88, which in turn releases IRAK^**62**^. Ultimately IRF7 is released which causes viral myocarditis. It is interesting to note that in human and mice, TLR7 is alternatively spliced and expressed as two protein isoforms^**63**^. Another interesting observation was that Chicken erythrocytes do not express TLR7^**64**^. While studying TLR7 expression pattern in different avian species, an interesting observation in our current study was that TLR 7 gene expression was significantly better in duck compared to other poultry species, as indigenous chicken breeds (Aseel, Haringhata Black chicken), goose and guineafowl. This is the first report for such a comparative study. It was reported that chicken TLR7 follow a restricted expression pattern. TLR7 expression was better in a macrophage cell line, chicken B-cell like cell line, but the expression was observed to be lower in kidney cell line^**65**^. Following Marek’s disease virus expression, TLR7 expression was observed to be increased in lungs^**66**^.

Similarly, increased TLR7 expression was noted in IBDV (infectious bursal disease virus)^**67**^. As regard to Avian influenza infection, it was observed that at the early stage of Low pathogenic avian influenza virus (LPAIV) infection of H11N9, both duck and chicken, TLR7 is transiently expressed in peripheral blood mononuclear cells (PBMC), while as infection progress, expression declines. Hence it was observed that in chicken TLR7 expression was depended on the interaction between host and RNA virus^**68**^. Thus differences in the expression pattern of TLR7 in chicken and duck was suggested^**68**^. Even in chicken, the TLR7 expression pattern was found to vary between HPAIV and LPAIV. Thus TLR7 was observed to be important immune response gene for avian influenza, TLR7 ligands show considerable potential for antivirals in chicken^**68**^. Although no direct report was available for better TLR7 expression in a duck in comparison to chicken, it was reported that tissue tropism and immune function of duck TLR7 is different from that of chicken TLR7, which result in a difference in susceptibility between chicken and duck, when infected by the same pathogen ^**68**^. A high expression pattern of duck TLR7 in respiratory and lymphoid tissue was observed to be different from that of chicken.

TLR3, TLR7, and TLR21 localize mainly in the ER in the steady-state and traffic to the endosome, where they engage with their ligands. The recognition triggers the downstream signal transduction to activate NF-κв or IRF3/7, finally induces interferon and inflammatory cytokine production^**68**^. We can explore these identified and characterized genes for production of transgenic or gene-edited chicken resistant to Avian influenza as a future control strategy against Avian influenza through immunomodulation, devoid of side effects as in case of use of drugs^**69**^. It is to be noted that since the control of Avian influenza virus has been difficult and challenging either through vaccination^**70,71,72**^ or treatment through antiviral drugs^**73,74,75,76**^due to frequent mutation and genetic reassortment (regarded as antigenic shift or antigenic drift) of the single stranded RNA genome which is prone to mutations^**77,78,79,80**^. An interesting observation revealed that unlike antibodies (comprising of immunoglobulins) which were highly specific, arising due to variability of Fab site and variable region^**81**^, immune response molecules for innate immunity can bind Avian influenza virus (H, N antigen), irrespective of strains. As we analyze the binding sites, some important domains were identified, which may be involved in antiviral activity. This led to the finding that therapeutic approach may be attempted with the recombinant product corresponding to identified domain. Gene editing with gene insert from identified gene may lead to the evolving of disease resistant strains/lines of chicken or duck.

Although the receptors for Human influenza and Avian influenza are different, mutations may overcome the barrier. As a case report in 2018, a human infection with a novel H7N4 avian influenza virus was reported in Jiangsu, China. Circulating avian H9N2 viruses were reported to be the origin of the H7N4 internal segments, unlike the human H5N1 and H7N9 viruses that both had H9N2 backbones. The major concern is that genetic reassortment and adaptive mutation of Avian influenza virus give rise to Human influenza virus strain H7N4 ^**82,83,84,85**^. WHO has also warned about pandemic on Human flu resulting from genetic reasssortment of Avian influenza^**86**^. We observed in this study that H5N1 strain of Avian influenza, a highly pathogenic strain was genetically closer to H6N2. Recent reports revealed that H6N2 is continuously evolving in different countries as South Africa^**87**^, Egypt^**88**^, India^**89**^, North America^**90**^ due to genetic reassortment. It is gradually evolving from low pathogenic form to high pathogenic form and observed to overcome species barrier with interspecies genetic assortment ^**91,92**^ and every possibility to evolve as pandemic for human. Reports are available depicting human influenza virus arising due to genetic reassortment of avian influenza in China^**82,83,84,85**^,. These findings highlights the growing importance of the study in current era, when the world is suffering from a pandemic.

Although molecular docking analysis were available for identification of various drug molecules with Avian influenza virus^**93,94,95**^, this is the first report of molecular docking analysis with the Immune response molecules responsible for antiviral immunity against Avian influenza and is the basis for finding drug for a disease. A series of immune response molecules are responsible for providing antiviral immunity with their respective interaction in various pathways as we depicted through String and KEGG pathway analysis in our current study.

## Conclusion

RIG1 detect the virus that is present within the cytosol of infected cells (cell intrinsic recognition), whereas TLR3 detects virus-infected cells, and TLR7 detects viral RNA that has taken up into the endosomes of sentinel cells (cell-extrinsic recognition).TLR7 may be regarded as the promising gene for antiviral immunity with pronounced expression profiling in duck in contrast to other poultry birds. Molecular docking revealed RIGI, TLR3 and TLR7 being the promising gene conferring antiviral immunity against Avian influenza. Point mutations have been detected in chicken TLR3 with respect to that of duck indicative of reduced antiviral immunity in chicken in comparison to duck.

## Acknowledgment

The authors are thankful to Department of Biotechnology, Ministry of Science and Technology, Govt. of India (Grant number BT/PR24310/NER/95/649/2017) and Department of Science and Technology, Govt. of India (Grant no. EMR/2016/003554) for providing the financial support. The technical and financial support by Vice-Chancellor, West Bengal University of Animal and Fishery Sciences is duly acknowledged. Thanks to Director, AH & VS, Animal Resource Development Department, Govt. of West Bengal.

## Competing interest

The author(s) declare no competing interests.

## Author’s contribution

Aruna Pal has designed the research work, conducted the research work, analyzed data and written the manuscript. Abantika Pal has conducted the bioinformatics analysis. PB has analyzed and revised the article.

## Notes

### Competing Interest Statement

The authors have declared no competing interest.

