## Supplementary figures and images for "Molecular characterization of RIGI, TLR7 and TLR3 as immune response gene of indigenous ducks in response to Avian influenza"

### Supplementary file 1

Supplementary Fig 1: PatchDock Analysis

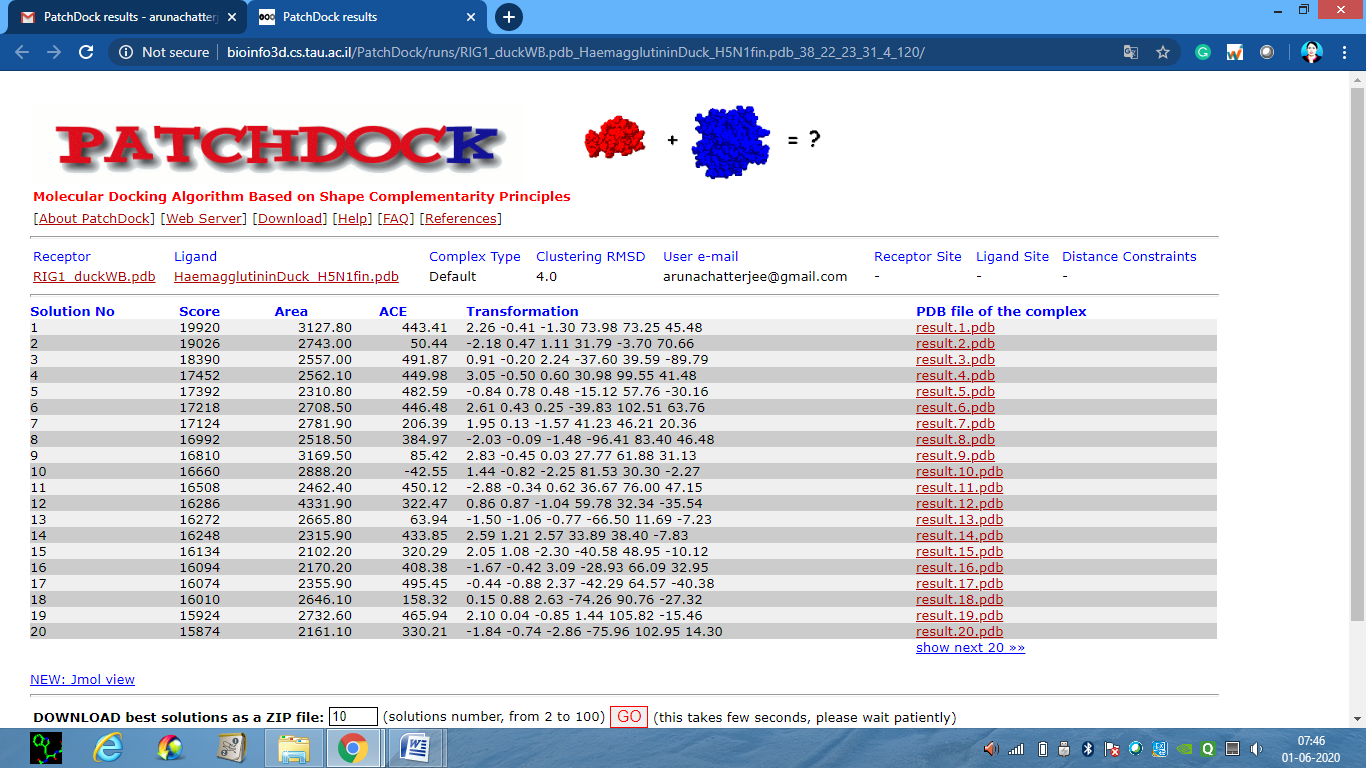

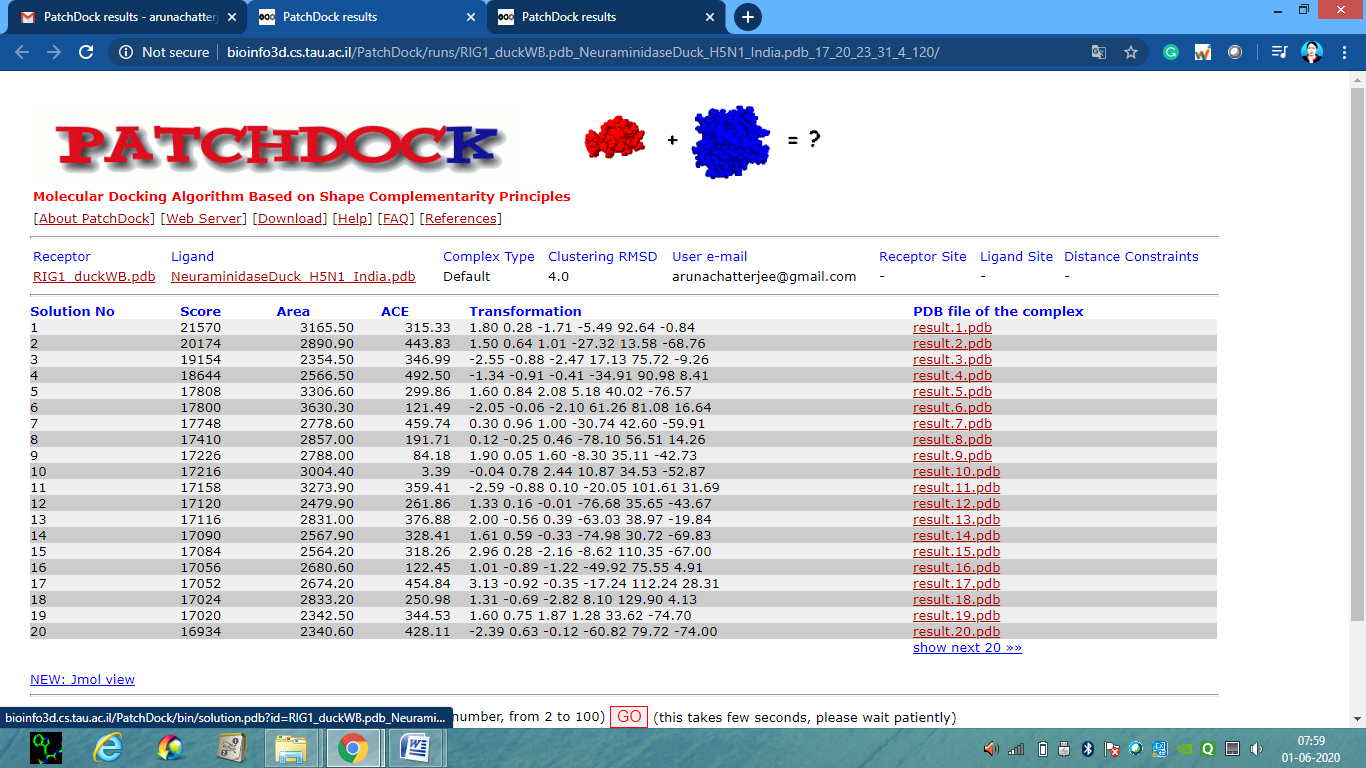

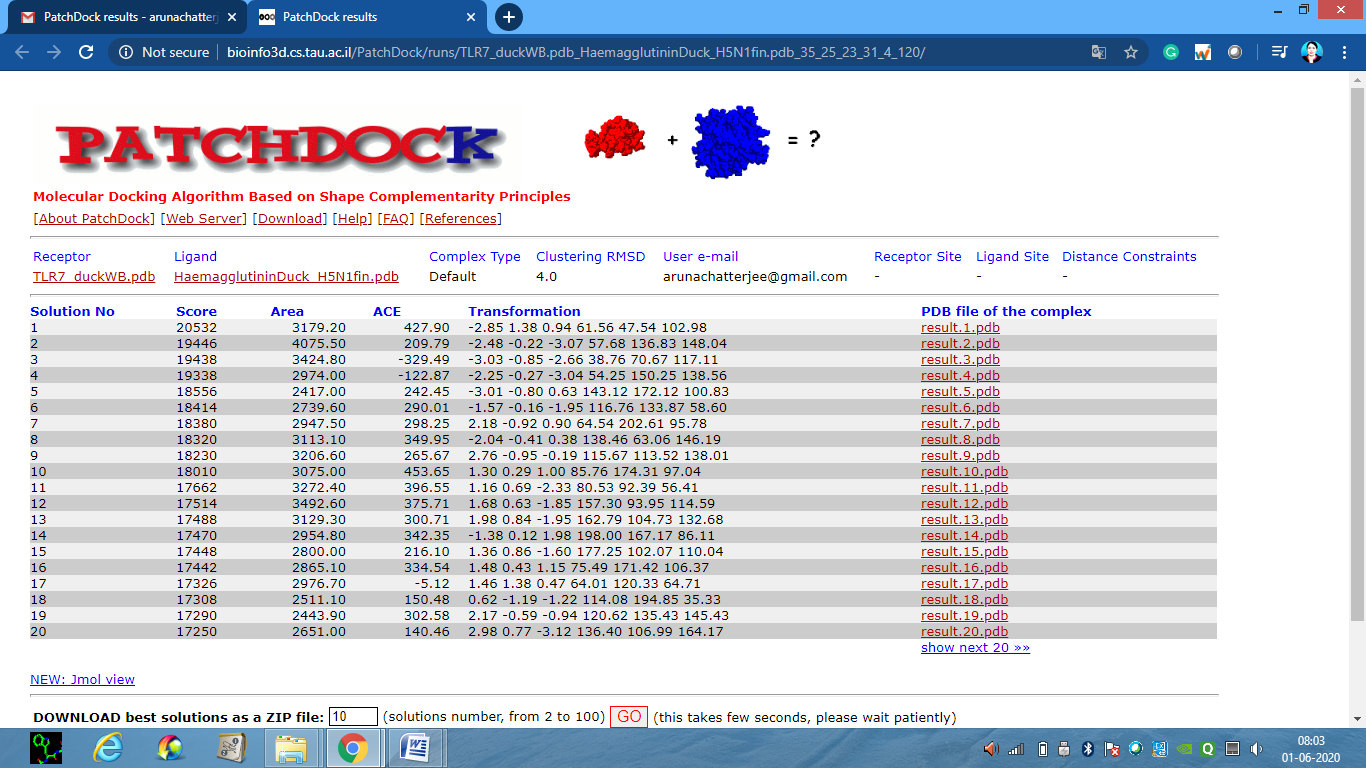

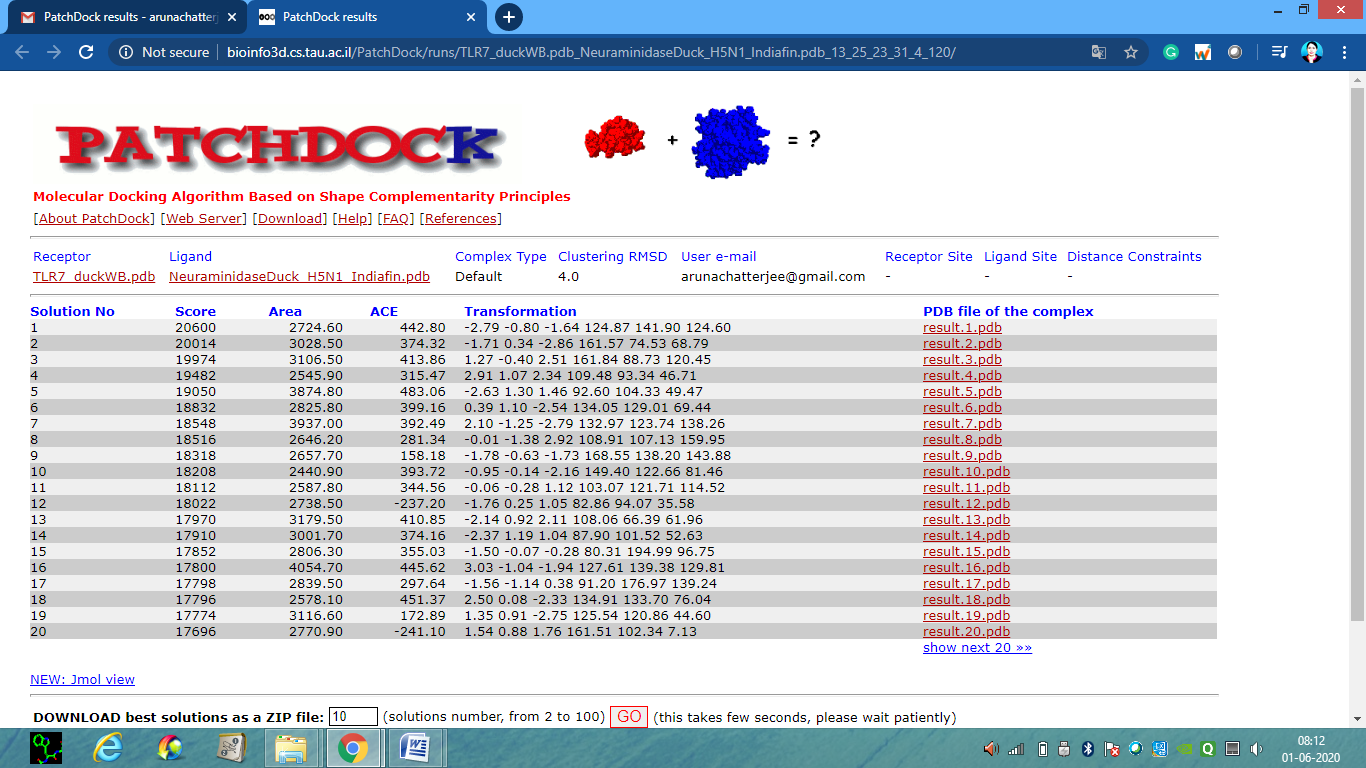

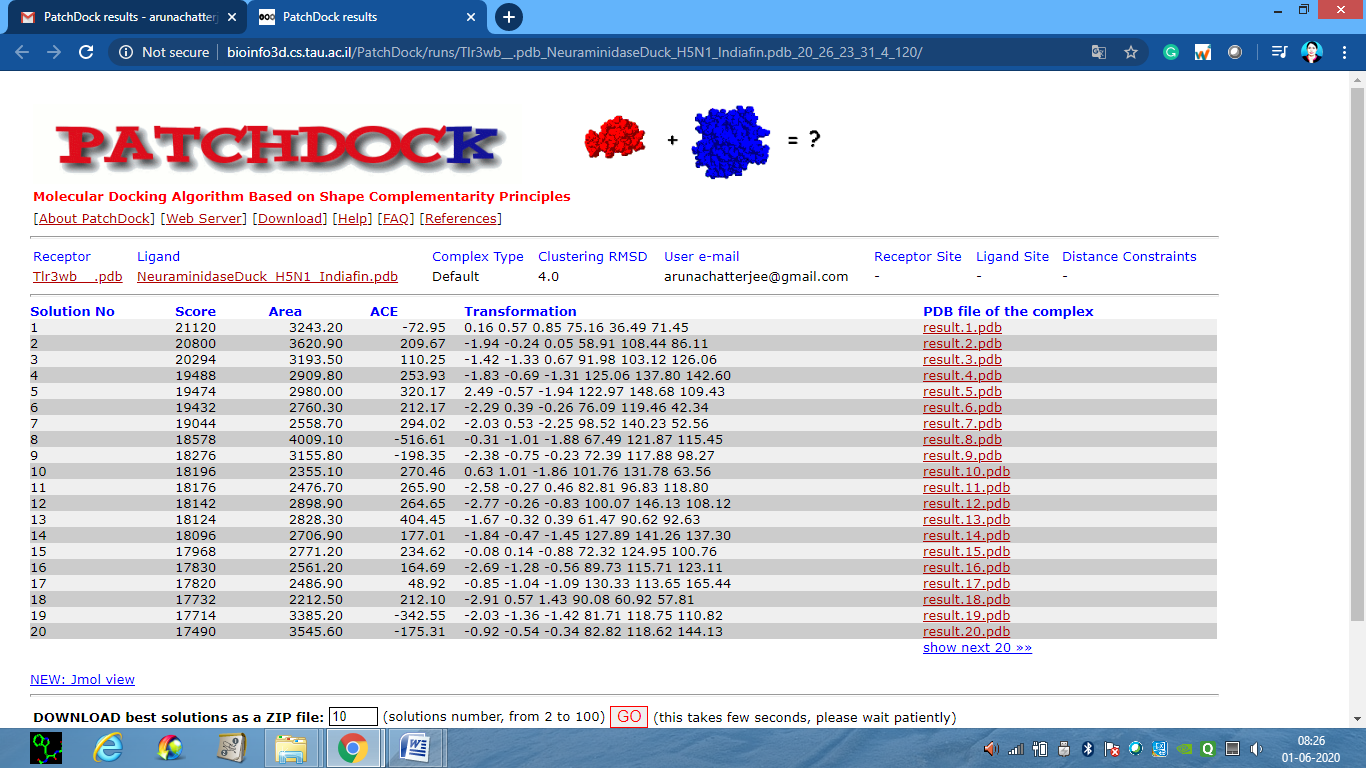

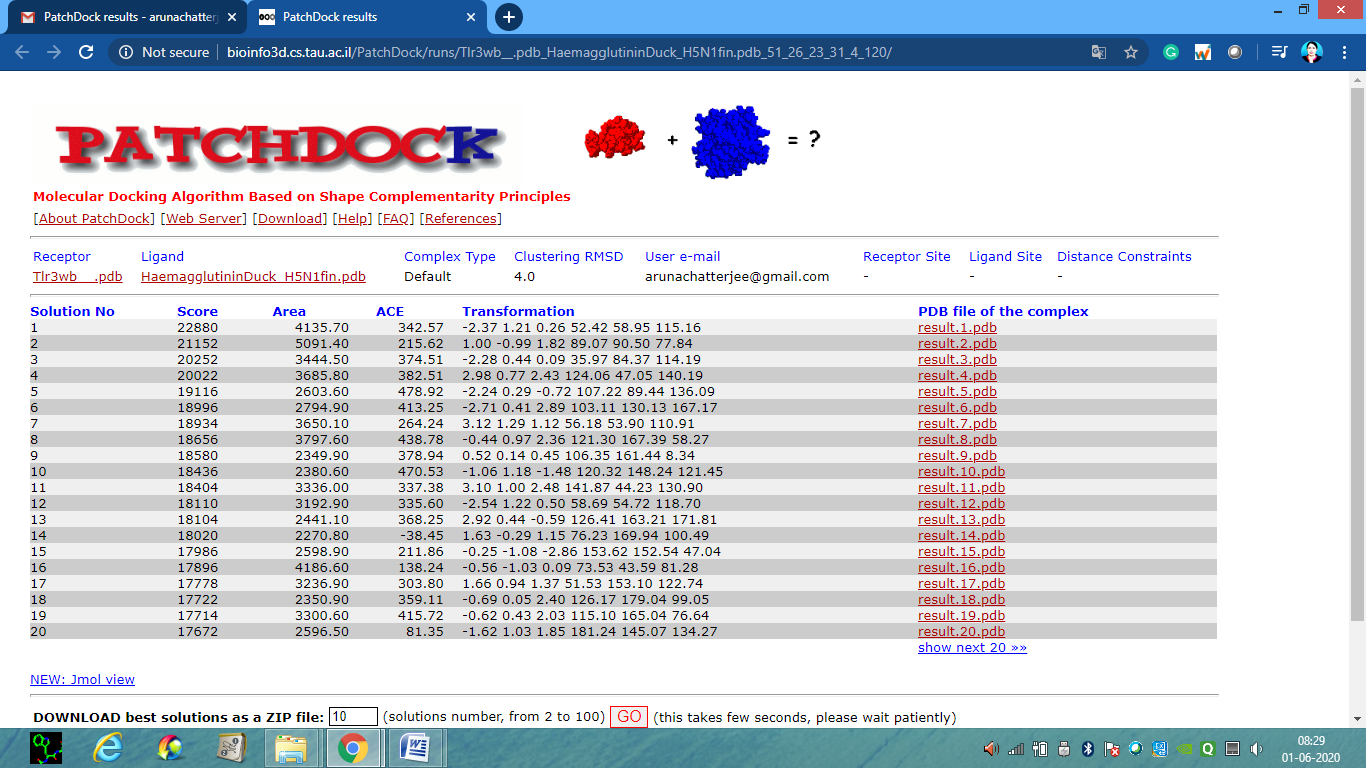

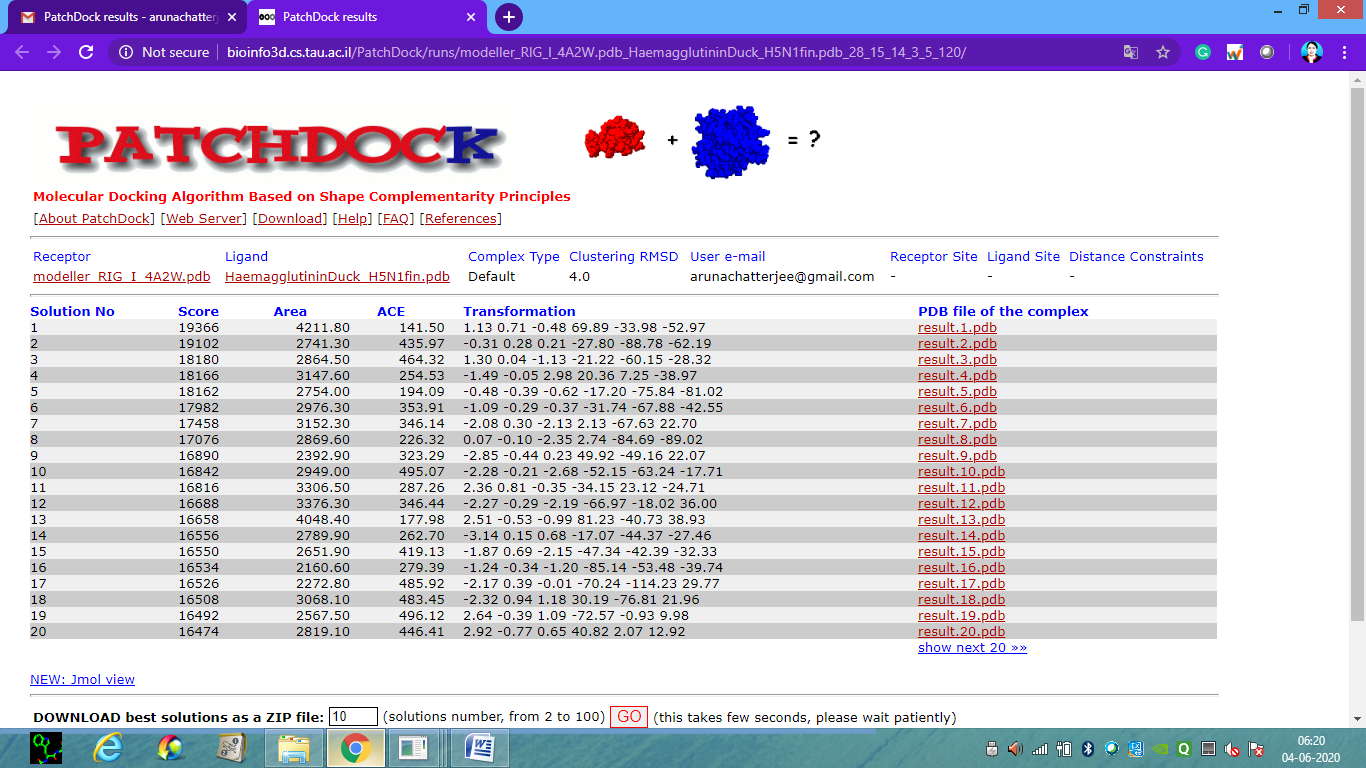

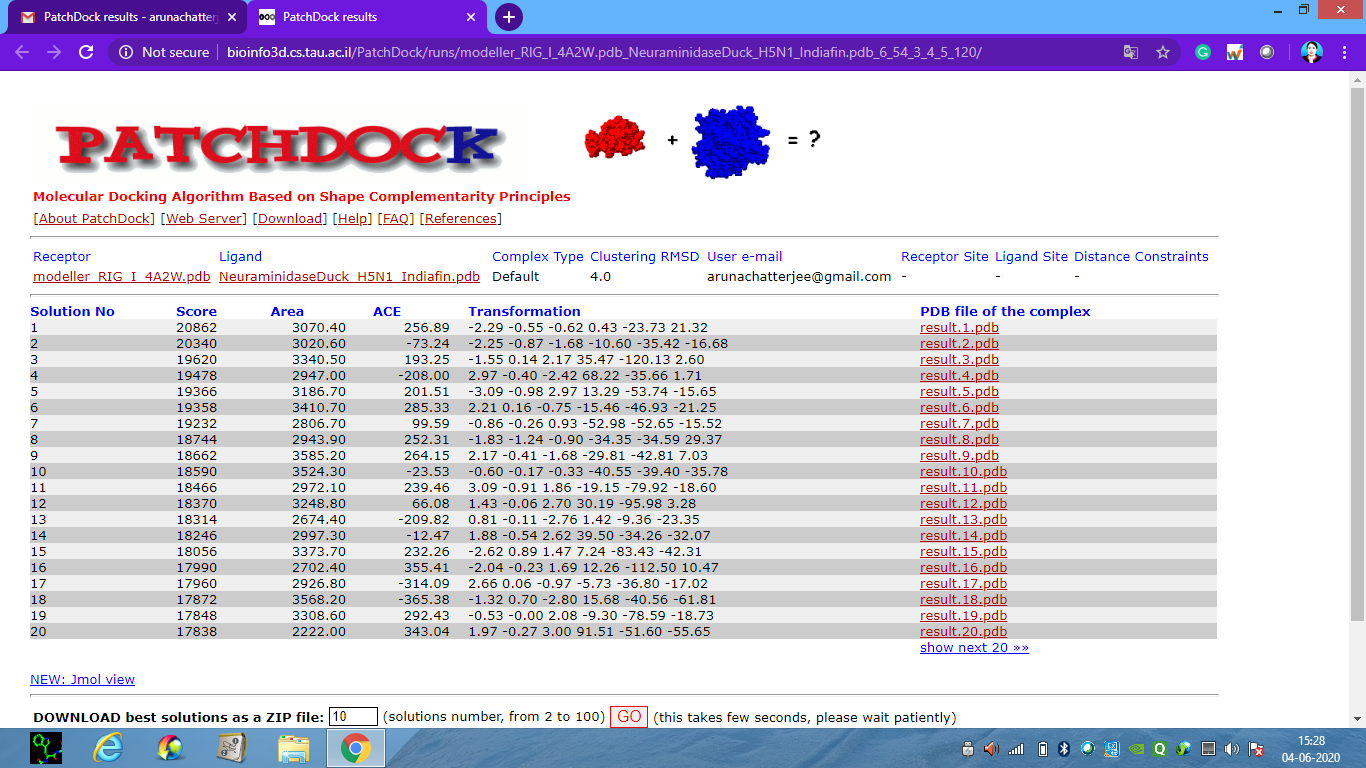
